# Potent and broadly neutralizing antibodies against sarbecoviruses induced by sequential COVID-19 vaccination

**DOI:** 10.1101/2023.08.22.554373

**Authors:** Xiaoyu Zhao, Tianyi Qiu, Xiner Huang, Qiyu Mao, Yajie Wang, Rui Qiao, Tiantian Mao, Yuan Wang, Jiayan Li, Cuiting Luo, Chaemin Yoon, Xun Wang, Chen Li, Yuchen Cui, Chaoyue Zhao, Minghui Li, Yanjia Chen, Guonan Cai, Wenye Geng, Zixin Hu, Jinglei Cao, Wenhong Zhang, Zhiwei Cao, Hin Chu, Lei Sun, Pengfei Wang

**Author notes:** These authors contributed equally (Xiaoyu Zhao, Tianyi Qiu, Xiner Huang, Qiyu Mao, Yajie Wang, Rui Qiao). Address correspondence to Pengfei Wang, Lei Sun, Hin Chu or Zhiwei Cao.

## Abstract

The current SARS-CoV-2 variants strikingly evade all authorized monoclonal antibodies and threaten the efficacy of serum-neutralizing activity elicited by vaccination or prior infection, urging the need to develop antivirals against SARS-CoV-2 and related sarbecoviruses. Here, we identified both potent and broadly neutralizing antibodies from a five-dose vaccinated donor who exhibited cross-reactive serum neutralizing activity against diverse coronaviruses. Through single B cell sorting and sequencing followed by a tailor-made computational pipeline, we successfully selected 86 antibodies with potential cross-neutralizing ability from 684 antibody sequences. Among them, one potently neutralized all SARS-CoV-2 variants that arose prior to Omicron BA.5, and the other three could broadly neutralize all current SARS-CoV-2 variants of concern, SARS-CoV and their related sarbecoviruses (Pangolin-GD, RaTG13, WIV-1, and SHC014). Cryo-EM analysis demonstrates that these antibodies have diverse neutralization mechanisms, such as disassembling spike trimers, or binding to RBM or SD1 to affect ACE2 binding. In addition, prophylactic administration of these antibodies significantly protects nasal turbinate and lung infections against BA.1, XBB.1 and SARS-CoV viral challenge in golden Syrian hamsters, respectively. This study reveals the potential utility of computational process to assist screening cross-reactive antibodies, as well as the potency of vaccine-induced broadly neutralizing antibodies against current SARS-CoV-2 variants and related sarbecoviruses, offering promising avenues for the development of broad therapeutic antibody drugs.

## Introduction

Severe acute respiratory syndrome coronavirus 2 (SARS-CoV-2), the causative agent of coronavirus disease 2019 (COVID-19), has caused over 769 million confirmed cases, along with more than 6.9 million deaths worldwide (https://covid19.who.int). During the course of COVID-19 pandemic, SARS-CoV-2 continuously evolves with changes in the genome caused by genetic mutations or viral recombination, resulting in variants that are different from the original SARS-CoV-2 virus^1–3^. The World Health Organization has now designated five SARS-CoV-2 variants of concern (VOCs), including Alpha (B.1.1.7), Beta (B.1.351), Gamma (P.1), Delta (B.1.617.2), and Omicron (B.1.1.529 or BA.1)^4^. In particular, the Omicron VOC contains an alarming number of over 30 mutations in its spike (S) protein and has spread rapidly worldwide^5^. This situation continues to evolve with increased global spreading of Omicron subvariants, including BA.2, BA.2.12.1, BA.2.75, BA.2.75.2, BA.4, BA.4.6, BA.5, BF.7, BN.1, BQ.1, BQ.1.1, CH.1.1, and the most recent XBB sublinages^6–9^.

Although COVID-19 vaccines and therapeutic antibodies have been developed and deployed at an unprecedented speed, the continued evolution of SARS-CoV-2 variants has raised concerns about the effectiveness of monoclonal antibody (mAb) therapies and the potential evasion from vaccine-induced immunity^10–12^. Recent studies have revealed that the efficacy of mRNA monovalent vaccine after two or three doses was less than 50% during the BA.4/5 waves, even a fourth dose only causing a minimal transient increase in Omicron-neutralizing antibodies^10, 13, 14^. In addition, BQ and XBB subvariants have been reported to have an immune evasion capacity higher than BA.5 in vaccinated patients^15–17^. We and other research groups have also shown that two doses of an inactivated whole-virion vaccine showed weak to no neutralization activity, although homologous or heterologous boosters improved neutralization titers against Omicron subvariants^4, 18, 19^. Moreover, several approved and clinical-stage mAbs suffer from a loss of efficacy against Omicron subvariants^20–22^. In particular, the Omicron subvariants BQ.1.1 and XBB.1.5 have resulted in marked or complete resistance to neutralization of almost all authorized antibodies^9^. Considering that the authorized or approved mAbs for clinical immunotherapy have shown greatly reduced activities, it is necessary to identify broadly neutralizing antibodies that fully cover the various SARS-CoV-2 variants.

Apart from SARS-CoV-2, SARS-CoV that use human angiotensin-converting enzyme 2 (ACE2) as receptor, and MERS-CoV belong to beta-coronaviruses using receptors other than ACE2, have previously posed a great threat to human health^23, 24^. In addition, the growing threat of continued zoonotic spillovers reveals the importance of the development of interventions that could broadly combat zoonotic coronaviruses with pandemic potential^25, 26^. For instance, a previous study reported that a SARS-CoV-2-related pangolin coronavirus exhibits similar infection characteristics to SARS-CoV-2 and can direct contact transmissibility in hamsters^27^. More recently, another pangolin-origin SARS-CoV-2-related coronavirus was also shown to have similar infectivity to SARS-CoV-2 in both human cells and organoids, highlighting the potential risk of spillover from pangolins and follow-up circulation in human populations^28^. Therefore, finding potent and broadly neutralizing mAbs that can target not only the circulating SARS-CoV-2 variants but also related sarbecoviruses is of utmost importance.

In this study, we reported the isolation of potent and broadly neutralizing mAbs from a vaccinated donor with a special five-dose COVID-19 vaccination schedule. One mAb, named PW5-570, displayed the most potent neutralizing activity against all SARS-CoV-2 strains prior to BA.5 variant, and another three mAbs, named PW5-4, PW5-5 and PW5-535 were found to broadly neutralize all sarbecoviruses tested, including all the current SARS-CoV-2 variants, SARS-CoV, Pangolin-GD, RaTG13, WIV-1, and SHC014. Structural analysis showed that these antibodies had a variety of characteristic binding modes, with PW5-5 and PW5-535 binding to different conserved epitopes hidden in RBD, while the epitope of PW5-570 overlapped with RBM to affect binding to ACE2. Moreover, prophylactic administration of PW5-570 fully protected nasal turbinate and lung infections against Omicron BA.1 viral challenge in golden Syrian hamsters, displaying promising biomedical interventions against pandemic SARS-CoV-2 VOCs. More importantly, PW5-5 and PW5-535 exhibited *in vivo* efficacy against both SARS-CoV-2 XBB.1 and SARS-CoV, suggesting their use as therapeutic agents for pan-sarbecoviruses antivirals. Taken together, from a single sequential vaccinated donor, we not only found an ultra-potent neutralizing mAb against SARS-CoV-2 VOCs, but also identified three broadly neutralizing mAbs with pan-sarbecoviruses potential. This study reveals the potency of vaccine-induced broadly neutralizing mAbs against current VOCs and other related sarbecoviruses, and affords the potential for broad therapeutic mAb drugs.

## Results

### Identification of a sequential vaccinated donor with cross-reactive serum neutralizing activity

To isolate potent broadly neutralizing mAbs against currently circulating SARS-CoV-2 variants and other human coronaviruses, we screened out one healthy volunteer who had received a total of five COVID-19 vaccination doses, starting with two doses of mRNA vaccine (BNT162b2) followed by three doses of inactivated whole-virion vaccine (BBIBP-CorV), as depicted in Figure 1A. We collected serum samples from this donor after the fourth and fifth vaccination, and tested their neutralizing antibody titer (ID_50_) against a panel of pseudotyped human coronaviruses, including SARS-CoV-2 WT (D614G), BA.1, BA.2, BQ.1.1, and XBB.1 variants, as well as SARS-CoV and MERS-CoV. For comparison, neutralization ID_50_ values were also measured for 11 local vaccinees, whose blood samples were collected at 1 month after the third BBIBP-CorV-vaccination (Figure 1B and Supplementary Figure 1). Although the Omicron subvariants showed substantial neutralization evasion from all the serum samples, similar as we reported previously^17^, serum from the sequential vaccination donor after the fourth and fifth dose had consistently higher neutralization titers than the three-dose inactivated vaccination group. Particularly, the serum after the fifth vaccination dose reached 3-13-fold higher neutralization titers than the mean of the three-dose inactivated vaccination group against SARS-CoV-2 and its variants. In addition, the serum after the fifth vaccination dose also showed 7-fold higher SARS-CoV neutralization titer than the mean of the three-dose inactivated vaccination group. More importantly, the four and five-dose sequential COVID-19 vaccination even elicited cross-reactive neutralizing antibodies against MERS-CoV, but not the three-dose homologous inactivated vaccination, which induced undetectable neutralization activity against MERS-CoV (ID_50_ <1:10) (Figure 1B). These results indicated that sequential immunization of heterologous COVID-19 vaccinations could elicit cross-reactive broadly neutralizing antibodies against SARS-CoV-2, its variants, SARS-CoV, and even MERS-CoV. We, therefore, chose this special vaccinee for subsequent search of broadly neutralizing mAbs.

**Figure 1.**
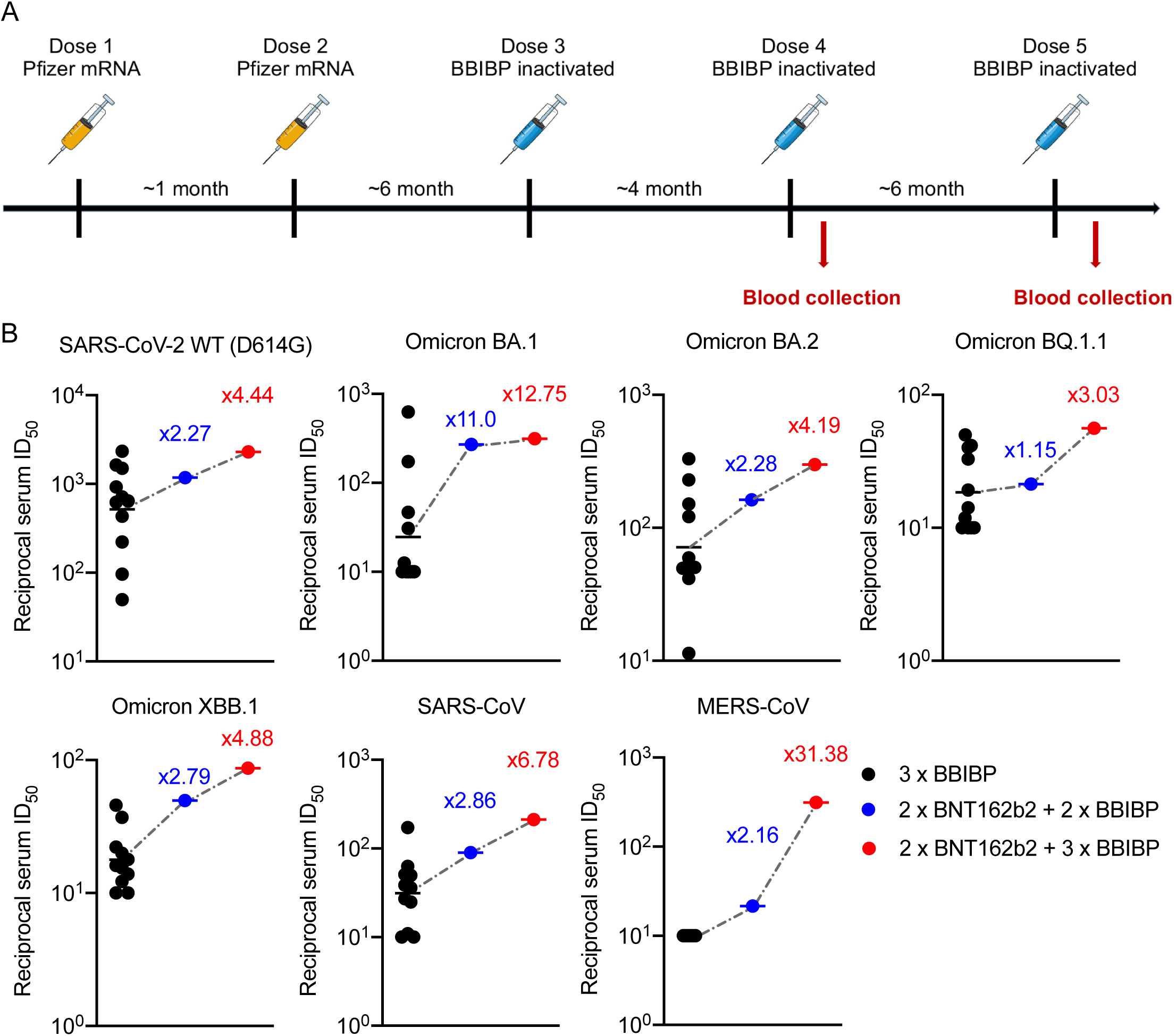
Cross-reactive immune responses elicited by sequential vaccination. **(A)** Vaccination schedule overview. A healthy volunteer who received a total of five doses of vaccine, starting with two-dose mRNA vaccine (BNT162b2) followed by three-dose inactivated whole-virion vaccine (BBIBP-CorV). Blood samples were collected after the 4^th^ and 5^th^ vaccination dose as illustrated in the diagram. **(B)** Neutralizing antibody titers of the sequential vaccination doner (4-dose: blue and 5-dose: red) comparing to the 3-dose BBIBP vaccinees (black, n = 11). Neutralizing antibody titers represent serum dilution required to achieve 50% virus neutralization (ID_50_). The numbers indicate the fold of enhancement of ID_50_ values relative to mean titer measured among 3-dose BBIBP vaccinees.

### Identification of broadly neutralizing antibodies against severe human coronaviruses

Peripheral blood mononuclear cell (PBMC) samples were collected from this selected donor 2 weeks after the fifth immunization dose. We then sorted for MERS-CoV or SARS-CoV-2 Omicron S trimer–specific memory B cells from the blood (Supplementary Figure 2), followed by single-cell RNA sequencing to determine the paired heavy and light chain sequences of each mAb. Subsequently, an *in-silico* screening was performed to identify potential cross-reactive mAbs for both MERS-CoV and SARS-CoV-2. Initially, the structure modeling of S protein from three coronaviruses (MERS-CoV, SARS-CoV-2 WT, and SARS-CoV-2 Omicron), each including four structure formats (1-RBD-up, 2-RBD-up, 3-RBD-up and 3-RBD-down) (Supplementary Figure 3), and the 684 antibodies were constructed. Then, we screened 1,083 real epitopes and 9,608 virtual epitopes of the coronavirus S proteins through CE-BLAST^29^, which found 132 potential cross-reactive epitopes. Finally, through the patch model of SEPPA-mAb^30^, we ranked the binding score between each cross-reactive epitope and the complementarity-determining region (CDR) of the antibodies through structure complementarity calculation. Eventually, 86 top-ranking mAbs for at least one cross-reactive epitope were isolated for further analysis (Figure 2A).

**Figure 2.**
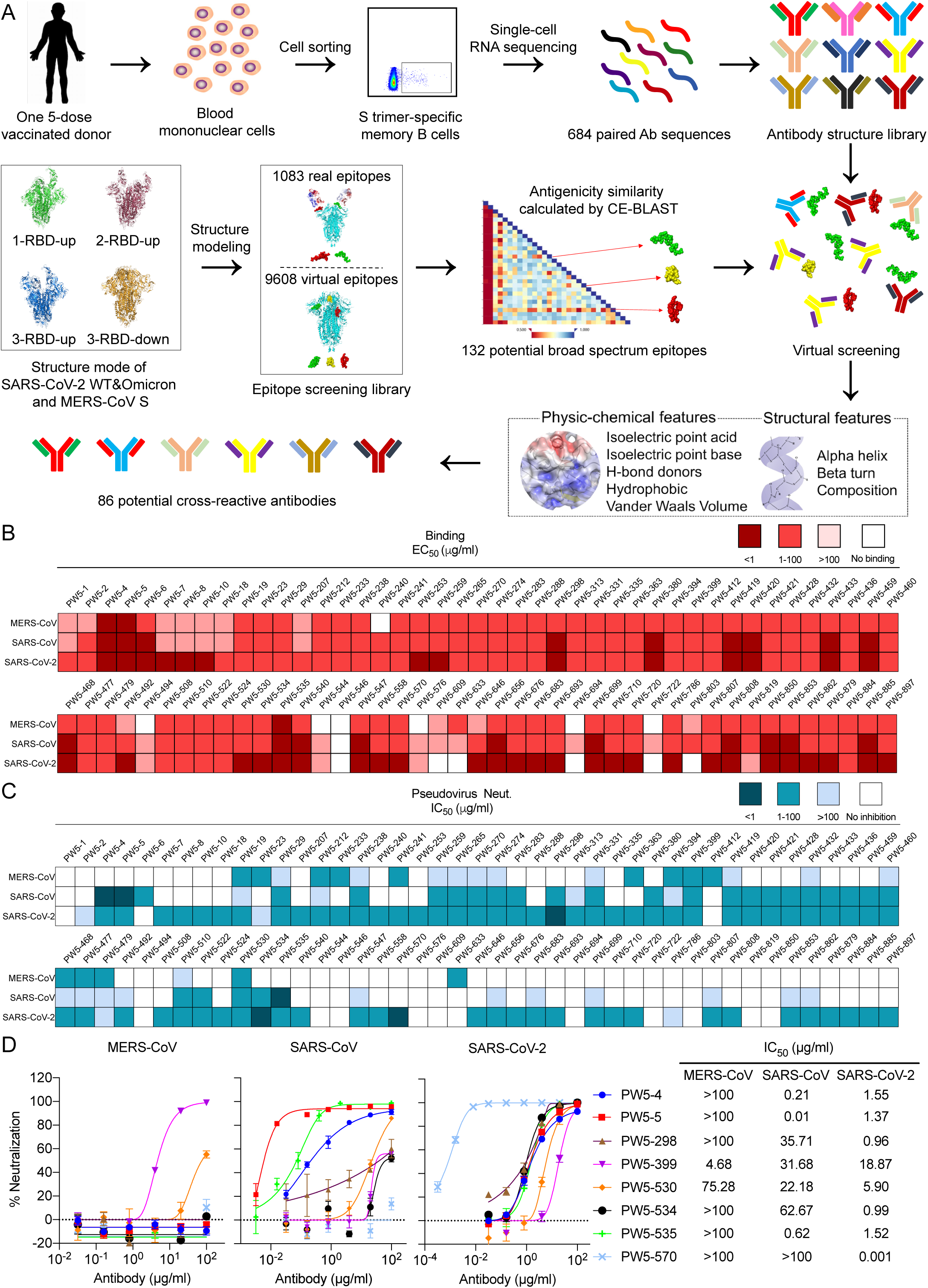
Identification of broadly neutralizing antibodies against severe human coronaviruses. **(A)** Schematic representation of the isolation process for potential broadly neutralizing antibodies using a combined method of single B cell sorting and virtual screening. **(B)** Heatmap showing the binding EC_50_ values of the selected mAbs with S trimers of MERS-CoV, SARS-CoV and SARS-CoV-2, respectively. **(C)** Heatmap showing the neutralization IC_50_ values of each mAb with indicated pseudotyped viruses. **(D)** Neutralization curves (Left) and IC_50_ values (Right) of indicated mAb with MERS-CoV, SARS-CoV and SARS-CoV-2 pseudotyped viruses, respectively. The data are representative of one of at least three independent experiments and are presented as the mean ± SEM.

To determine whether there were differences between the selected 86 mAbs and the total 684 mAbs, we characterized their heavy and light chain V gene usage, CDR3 length, and the number of nucleotide mutations. The results showed that apart from the same V gene usage, IGHV3-11, IGHV3-21, IGKV2D-28, IGKV 2-30, IGLV3-1, IGLV2-11 and IGLV3-25 were enriched in the selected 86 mAbs compared with the 684 mAbs (Supplementary Figure 4A and 4B). For the CDR3 length, no significant difference was observed between the 86 mAbs and the 684 mAbs, and the peak of the heavy chain and light chain is about 16 aa and 11 aa, respectively (Supplementary Figure 4C and 4D). Notably, number of nucleotide mutations in the 86 mAbs is significantly higher than that in the 684 mAbs, indicating that the selected 86 mAbs have a more violent mutation than the rest of antibodies in the 684 mAbs (Supplementary Figure 4E and 4F).

To determine the cross-binding abilities of the 86 mAbs, we assayed the binding EC_50_ values of these mAbs to the S trimer of MERS-CoV, SARS-CoV and SARS-CoV-2, respectively. We found that most of the mAbs to be highly cross-reactive to the S trimers of SARS-CoV and SARS-CoV-2 but not to that of MERS-CoV, whereas PW5-4, PW5-5 and PW5-535 displayed high binding ability to all of them (Figure 2B). We further assayed the cross-activities of those mAbs against MERS-CoV, SARS-CoV and SARS-CoV-2 by VSV-based pseudotyped virus neutralization. Consistent with the binding profiles for these mAbs, most of the mAbs showed cross-neutralizing activity to SARS-CoV and SARS-CoV-2 but less to MERS-CoV (Figure 2C). Among them, PW5-4, PW5-5, PW5-298, PW5-534, and PW5-535 showed good neutralizing activity against both SARS-CoV and SARS-CoV-2, while PW5-399 and PW5-530 displayed better broadly neutralizing potential, covering all three severe human coronaviruses including MERS-CoV. Another mAb PW5-570, although only showed neutralizing activity against SARS-CoV-2, its activity was ultra-potent with IC_50_of 1 ng/ml. The pseudovirus neutralization profiles for these selected 8 mAbs were shown in Figure 2D. These results demonstrated that ultra-potent SARS-CoV-2 neutralizing antibodies, as well as broadly neutralizing antibodies against other human coronaviruses, could be elicited by sequential SARS-CoV-2 vaccination.

### Characterizing the potency and breadth of neutralization conferred by selected antibodies

To understand the breadth of these 8 newly cloned mAbs, we first performed neutralization assays using pseudotyped SARS-CoV-2 VOC viruses, including Alpha, Beta, Gamma, Delta, and Omicron (BA.1) (Figure 3A and 3E). All mAbs, except PW5-399 and PW5-530, showed neutralization breadth against all SARS-CoV-2 VOCs. PW5-570 was the most potent with extremely low IC_50_ values, followed by a group composed of PW5-4, PW5-5, PW5-298, PW5-534, and PW5-535. We next evaluated each mAb for neutralization against major SARS-CoV-2 Omicron subvariants, including BA.2, BA.5, BQ.1.1 and XBB, as well as its latest subvariants, such as XBB.1, XBB.1.16, XBB.2.3.3, EG.5.1, EU.1.1 and FY.4 (Figure 3B and 3E). Only PW5-4, PW5-5 and PW5-535 neutralized all Omicron subvariants tested, whereas other mAbs showed reduced or abolished neutralization activity for certain viruses in the panel. In addition, we evaluated each mAb for neutralization against four SARS-CoV or SARS-CoV-2 related sarbecoviruses that are capable of using human ACE2 as receptor. We found that all mAbs, except PW5-298, neutralized the two SARS-CoV-2 related sarbecoviruses (Pangolin-GD and RaTG13), and PW5-570 displayed the most potent neutralizing activity (Figure 3C and 3E). In contrast, for SARS-CoV related sarbecoviruses (WIV1 and SHC014), only PW5-4, PW5-5 and PW5-535 retained neutralizing activity, but not the other mAbs (Figure 3D and 3E).

**Figure 3.**
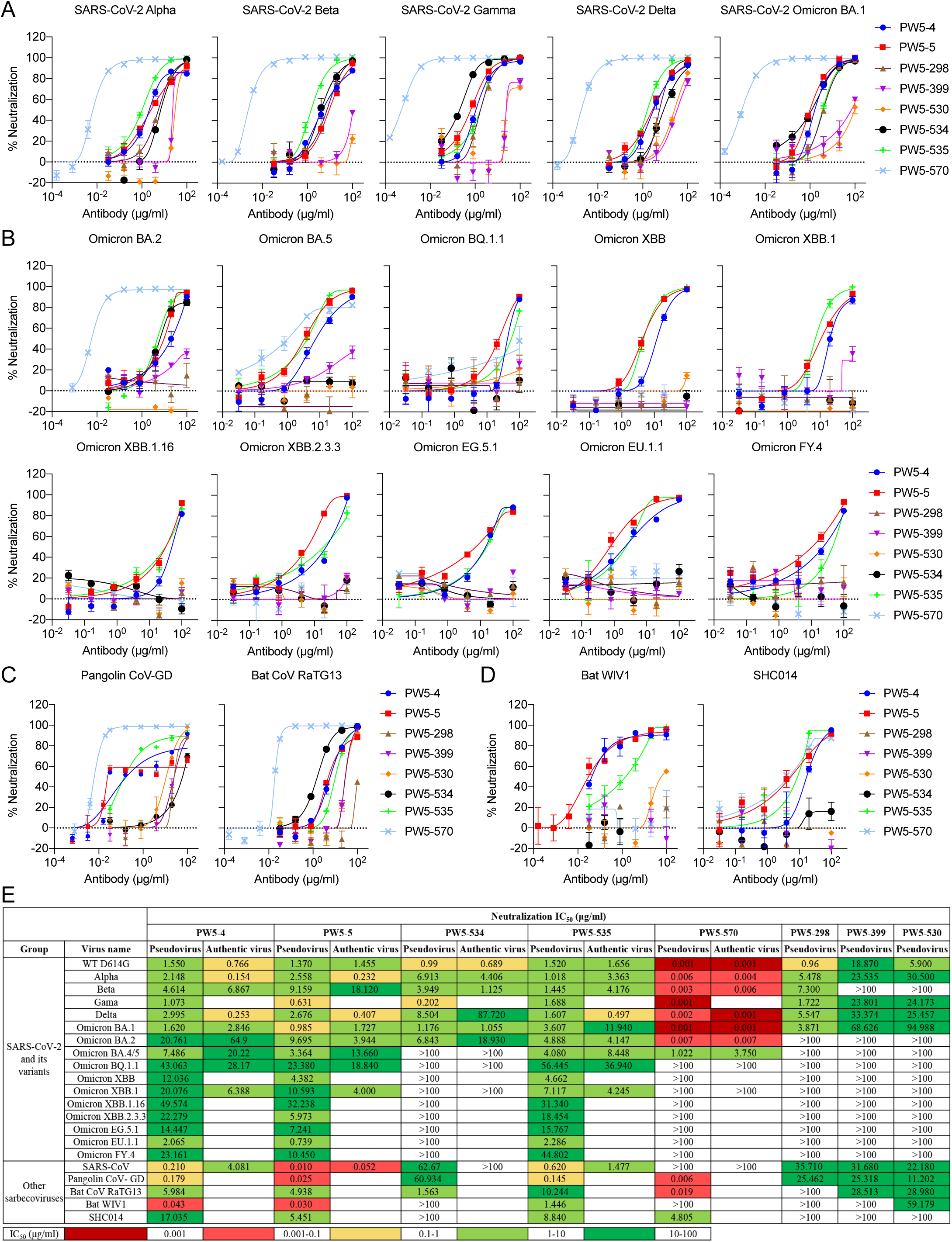
Neutralization of pseudotyped and authentic SARS-CoV-2 variants, SARS-CoV and other sarbecoviruses. **(A)** Neutralization of SARS-CoV-2 variants of concern (Alpha, Beta, Gamma, Delta, and Omicron BA.1) by indicated mAb. **(B)** Neutralization of major SARS-CoV-2 Omicron subvariants, including BA.2, BA.5, BQ.1.1 and XBB, as well as its latest subvariants, such as XBB.1, XBB.1.16, XBB.2.3.3, EG.5.1, EU.1.1 a nd FY.4 by indicated mAb. **(C)** Neutralization of SARS-CoV-2 related sarbecoviruses (Pangolin CoV-GD and Bat CoV RaTG13) by indicated mAb. **(D)** Neutralization of SARS-CoV related sarbecoviruses (Bat WIV1 and SHC014) by indicated mAb. **(E)** IC_50_ neutralization values of each mAb against SARS-CoV, SARS-CoV-2 and the major SARS-CoV-2 variant pseudoviruses as well as the live viruses. The data are representative of one of at least three independent experiments and are presented as the mean ± SEM.

We next performed authentic virus neutralization assays in Vero-E6 cells for five selected mAbs (PW5-4, PW5-5, PW5-534, PW5-535 and PW5-570) with better neutralization activity (Figure 3E). Similar as the pseudovirus results, we consistently found that PW5-4, PW5-5 and PW5-535 showed breadth by neutralizing all tested SARS-CoV and SARS-CoV-2 variants. Meanwhile, PW5-570 was the most potent mAb for all SARS-CoV-2 strains prior to the Omicron BA.5 variant. Besides, the neutralizing titers against the pseudoviruses correlated quite well with the titers obtained against the live virus (Supplementary Figure 5). Taken together, these results demonstrated that PW5-570 potently neutralized all SARS-CoV-2 strains prior to the Omicron BA.5 variant, but with no cross-neutralizing activity against SARS-CoV. While PW5-4, PW5-5 and PW5-535 neutralized all the tested SARS-CoV-2 variants as well as SARS-CoV and other related sarbecoviruses, PW5-5 exhibited better potency than the other two.

To further characterize these selected antibodies and investigate the potential for antibody combination, we measured the binding affinity and examined the competition between the mAbs and four other well-defined mAbs (CB6, LY-CoV555, S309 and CR3022) using the enzyme linked immunosorbent assay (ELISA)^31–34^. We found that all the five selected antibodies had a high affinity for the receptor-binding domain (RBD) but not the N-terminal domain (NTD) of the S protein (Supplementary Figure 6A). Besides, the competition profile indicated both overlapping and distinct epitopes recognized by these antibodies (Supplementary Figure 6B). PW5-4 and PW5-5 with overlapping epitopes, as well as the PW5-535, competed with the CR3022 (Class IV) for RBD binding. This analysis additionally revealed that through steric hindrance or binding-induced conformational change in RBD, PW5-570 is at least partially competitive with both CB6 (Class I) and LY-CoV555 (Class II), while PW5-534 is competitive with both LY-CoV555 (Class II) and S309 (Class III). We next performed a competition assay on S binding between the antibodies and ACE2 to investigate the neutralizing mechanism of these antibodies. As shown in Supplementary Figure 6C and 6D, the binding of ACE2 to SARS-CoV-2 S trimer was suppressed when PW5-534, PW5-535 and PW5-570 were mixed with SARS-CoV-2 S trimer, indicating a competition between these antibodies and ACE2. Thus, these results suggested that sequential vaccination could induce a broad spectrum of antibodies with distinct epitopes similar to those following natural infection.

### Cryo-EM structure of PW5-570 complexed with Omicron BA.1 S trimer

To understand the mechanism of SARS-CoV-2 Omicron variants potently neutralized by PW5-570, we determined the structure of Omicron BA.1 S trimer in complex with PW5-570 using cryo-electron microscopy (cryo-EM) (Figure 4A). The purified BA.1 S trimer was mixed with PW5-570 at a 1:1.2 molar ratio, incubated at 4°C for 1 h, and further purified by gel filtration. The peak fraction of the gel filtration was used for cryo-EM data collection (Supplementary Figure 7A). Unsymmetrical trimer dimers cross-linked by Fc regions were observed in 2D classification (Supplementary Figure 13). S trimers with 3 RBDs up, each binding to one IgG, were solved to 2.78Å resolution, and the S RBD with PW5-570 Fab region was locally refined to 2.93Å.

**Figure 4.**
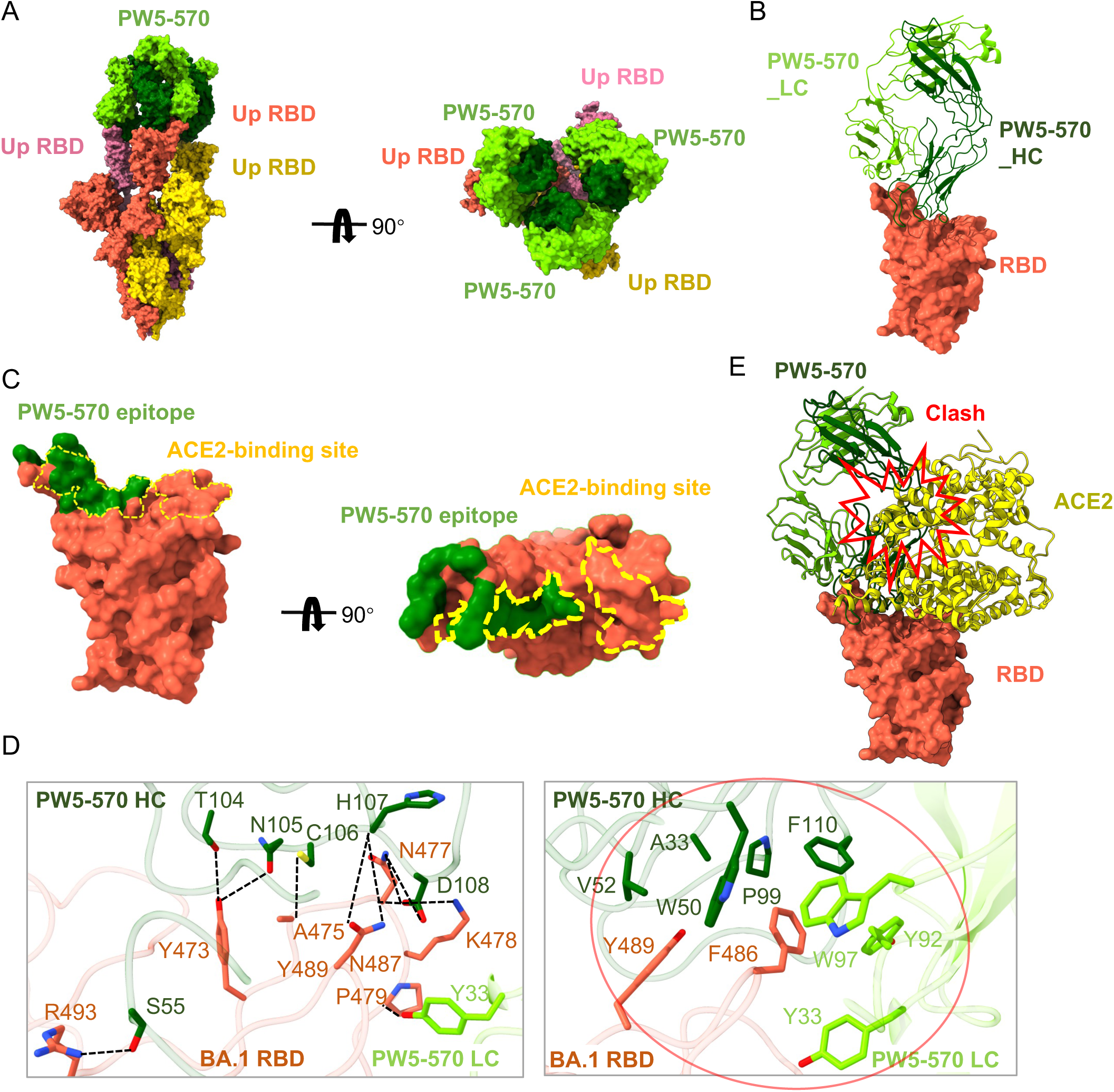
Cryo-EM structure of PW5-570 with Omicron BA.1 S trimer. (**A**) Cryo-EM structures of the BA.1 S trimer in complex with the antibody PW5-570 IgG. (**B**) Structure of BA.1 RBD-PW5-570. The RBD is displayed in tomato surface mode. The heavy chain and light chain of PW5-570 are shown as ribbons colored in dark green and light green, respectively. (**C**) Close-up view of the PW5-570 epitope on the RBD. ACE2 binding epitopes are represented by yellow dashed lines. (**D**) Detailed interactions between the PW5-570 and BA.1 S RBD. Hydrogen bonds are represented by dashed lines. Hydrophobic interactions are marked with red circles. (**E**) PW5-570 clash with the ACE2, ACE2 is marked yellow.

Structure analysis shows that the epitope of PW5-570 mostly overlaps with the RBM (Figure 4B and 4C). The binding of PW5-570 to RBD covers a surface area of 793 Å^2^ and involves a total of 12 RBD residues (Figure 4D). S55 of CDRH2 and T104, N105, C106, H107, D108 of CDRH3 contacts R493, Y473, A475, N487, N477 and K478 through 9 pairs of hydrogen bonds, and Y33 of CDRL1 forms a hydrogen bond with P479. In addition, A33 of CDRH1, W50, V52 of CDRH2, P99, F110 of CDRH3, Y33 of CDRL1, Y92, and W97 of CDRL3 form a hydrophobic interface with F486 and Y489. The binding epitopes of PW5-570 to RBD include the key residues Y473, A475, F486, and N487, required for ACE2 recognition and binding. Superposing the structures of the RBD/PW5-570 and RBD/ACE2 complexes showed extensive clashes between the antibody and the ACE2 receptor (Figure 4E), suggesting that binding of PW5-570 to RBD produces steric hindrance and prevents S from binding to the receptor ACE2. This agrees with the RBD-ACE2 blocking capacity of PW5-570 (Supplementary Figure 6C and 6D).

To further verify the key mutations conferring PW5-570 resistance, we constructed pseudoviruses with each of the single mutations alone or in combination based on Omicron BA.2 S. Consistent with cryo-EM structure of PW5-570, F486V is indeed the key mutation responsible for the loss in potency of PW5-570, while R493Q have the effect of mitigating its escape (Supplementary Figure 8).

### Cryo-EM structure of PW5-5 complexed with Omicron XBB and SARS-CoV S trimer

To understand the broad neutralization mechanism of PW5-5, we determined the cryo-EM structure of PW5-5 complexed with the stabilized prefusion ectodomain of Omicron XBB S (6P) and SARS-CoV S (2P), respectively. The purified XBB S trimer was mixed with PW5-5 at a 1:1.2 molar ratio, incubated at 4°C for 1 h, and further purified by gel filtration. The peak fraction of the gel filtration was used for negative-staining EM imaging, showing that PW5-5 binding disassembled XBB S trimer (Supplementary Figure 9). Therefore, we shortened the incubation time to 15 min and frozen the sample directly without further purification. The same strategy was used for SARS-CoV S-PW5-5.

Cryo-EM characterization revealed three conformational states of XBB S-PW5-5 complex: the XBB S trimer with two up-RBDs binding two PW5-5 (state 1), the XBB S trimer with three up-RBDs binding two PW5-5 (state 2), and the XBB S monomer with one PW5-5 (state 3), with resolution ranging from 3.1Å to 3.7Å (Figure 5A). Thus, binding of IgG PW5-5 results in S trimer disassembly. The structure of monomer XBB S-PW5-5 was at sufficient resolution for model building. PW5-5 binds to a covert epitope inside the RBD, burying a 936 Å^2^ surface area (Figure 5B and 5C). Totally 16 RBD residues are involved in the interaction. Among these, D54 of CDRH2 and R91 of CDRL3 from salt bridges with R357 and D428 of RBD, respectively. Y33 of CDRH1, N52, D54, S57, T59 of CDRH2, F102 of CDRH3, Y32 of CDRL1, Y49 of CDRL2 and N93 of CDRL3 form 10 pairs of hydrogen bonds with E516, Y396, H519, P463, D427, D428 and K462 (Figure 5D). Additionally, Y33, V50, V58 of CDRH2, F101, F102, P104, A105 of CDRH3 and W94 of CDRL3 form a hydrophobic patch with F464, P426, P463, L517, L518, A520 and P521 of RBD. In SARS-CoV S-PW5-5 complex, two confirmations were observed, the monomer state and the 2 up-RBDs, 2 Fabs state (Supplementary Figure 10A). The S RBD-PW5-5 Fab region was locally refined to 3.04Å resolution. PW5-5 interacts with SARS-CoV RBD in a similar way, burying 978 Å^2^ surface area (Figure 5E).

**Figure 5.**
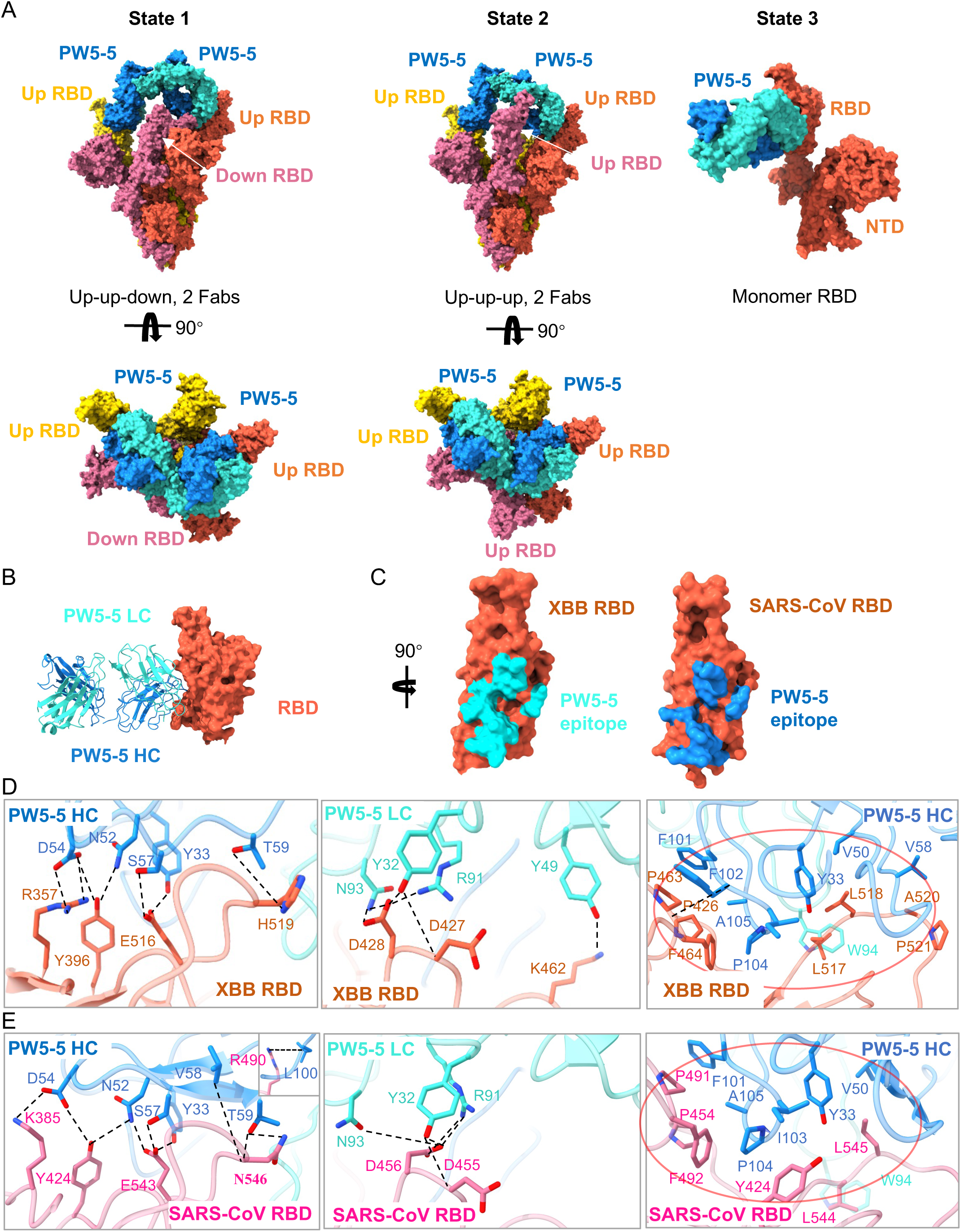
Cryo-EM structure of PW5-5 with Omicron XBB and SARS-CoV S trimer. (**A**) Cryo-EM structures of the XBB S in complex with the antibody PW5-5 IgG. (**B**) Structure of XBB RBD-PW5-5. The RBD is displayed in tomato. The heavy chain and light chain of PW5-5 are shown as ribbons colored in dark blue and light blue, respectively. (**C**) Close-up view of the PW5-535 epitope on XBB and SARS-CoV RBD. (**D**) Detailed interactions between the PW5-5 and XBB S RBD. (**E**) Detailed interactions between the PW5-5 and SARS-CoV S RBD. Hydrogen bonds are represented by dashed lines. Hydrophobic interactions are marked with red circles.

PW5-5 binding induces an large movement of the up-RBD: an outward rotation of 43.3° (up-RBD 1 in the 3-up state) and 55.9° (up-RBD 2 in the 3-up state) compared to the up-RBD in apo Omicron S (PDBID: 7WVN). This rotation is extraordinarily wider than previously reported antibodies FD01 (PDBID: 7WOQ) and bn03 (PDBID: 7WHK) with similar epitopes (Supplementary Figure 10B)^35, 36^. Such large deformation on S RBD may lead to instability. As no complex of SARS-CoV S trimer with 3 PW5-5 IgGs with was observed, we propose that binding of the third IgG result in the disassembly of the S trimer. In summary, the binding of PW5-5 induces RBD into an extra-wide-open state, resulting in S trimer instability and disassembly.

### Cryo-EM structure of PW5-535 complexed with Omicron XBB and SARS-CoV S trimer

We also determined the structures of PW5-535 complexed with the stabilized prefusion ectodomain of Omicron XBB S (6P) and SARS-CoV S (2P). The same purification strategy as BA.1-PW5-570 was used for XBB S-PW5-535 and SARS-CoV PW5-535 (Supplementary Figure 7B and 7C). Two states of XBB S-PW5-535 particles were observed, apo XBB S and 3 up-RBDs with 3 PW5-535 (Figure 6A). The cryo-EM structures of 3 up-RBDs state were determined to 2.95Å, with the RBD and PW5-535 region locally refined to 3.40 Å. As a contrast, two states of SARS-CoV S-PW5-535 were observed: 3 up-RBDs with 3 PW5-535 (3.05Å), and RBD monomer with PW5-535 (3.54Å) (Figure 6B). Notably, monomer state is only observed in SARS-CoV S-PW5-535 but not in XBB S-PW5-535, implying that SARS-CoV S(2P) itself is not as stable as XBB S (6P).

**Figure 6.**
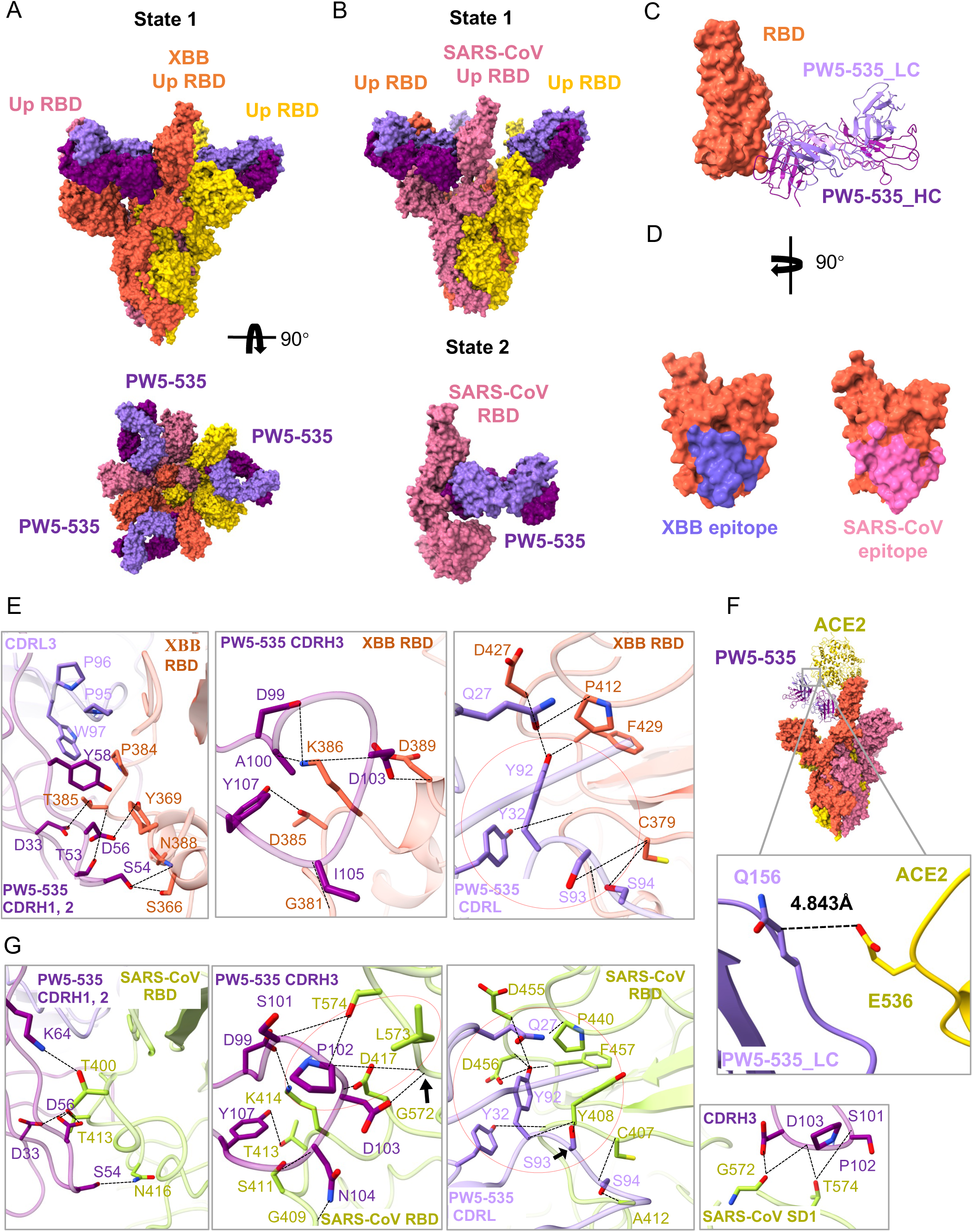
Cryo-EM structure of PW5-535 with Omicron XBB and SARS-CoV S trimer. (**A**) Cryo-EM structures of the XBB S in complex with the antibody PW5-535 Fab. (**B**) Cryo-EM structures of the SARS-CoV S in complex with the antibody PW5-535 Fab. (**C**) Structure of XBB RBD-PW5-535. The RBD is displayed in tomato. The heavy chain and light chain of PW5-535 are shown as ribbons colored in dark purple and light purple, respectively. (**D**) Close-up view of the PW5-535 epitope on the XBB RBD and SARS-CoV RBD. (**E**) Detailed interactions between the PW5-535 and XBB S RBD. (**F**) PW5-535 causes steric hindrance with ACE2. (**G**) Detailed interactions between the PW5-535 and SARS-CoV S RBD and SD1. Hydrogen bonds are represented by dashed lines. Hydrophobic interactions are marked with red circles.

In XBB S-PW5-535 complex, PW5-535 binds to a hidden epitope in RBD outside the RBM, buries a surface area of 961 Å^2^ and involves a total of 17 RBD residues (Figure 6C, 6D and 6E). Among these RBD residues, K386 is engaged in salt bridges with D99 and D103 of CDRH3, T385, S366, N388, Y369, K386, D389, G381, P412, D427, F429, C379 and V382 form 16 hydrogen bonds with residues from CDRs of PW5-535. Besides, Y58 of CDRH2, P95, P96, W97 of CDRL3 forms hydrophobic interaction with Y369 and P384. The R30 of CDRH1 interacts with the T531 of SD1 by the van der Waals force. SD1 is a subdomain closely related to RBD and highly conserved, which also explains that most antibodies against SD1 have a broad neutralization effect. In addition, the structural superimposition of Omicron S RBD-ACE2 (PDBID: 7WVP) onto XBB S timer-PW5-535 reveals a close contact between PW5-535 and ACE2, with only 4.8Å between Q156 of PW5-535 light chain and E356 of ACE2 (Figure 6F), indicating that PW5-535 binding would introduce a steric hindrance and prevent the virus from receptor recognition. This explains why PW5-535 competitively blocked the binding of ACE2 to SARS-CoV-2 S (Supplementary Figure 6C and 6D). PW5-535 induces SARS-CoV S disassembly, and buries a larger surface area of 1128 Å^2^ on SARS-CoV S monomer, involving 22 RBD residues, generating 24 pairs of hydrogen bonds and forming 2 patches of hydrophobic interactions. Compared with Omicron XBB, slight differences are observed in the interaction between PW5-535 and SARS-CoV S monomer (Figure 6G). Firstly, K64, P102, N104 of heavy chain forms extra hydrogen bonds with T400, G572, T574, L573, G409 and S411. Secondly, Y58 of heavy chain and P85, P96, and W97 of light chain don’t form hydrophobic interaction but replaced by a hydrophobic patch between P102 of CDRH3 and L573. Thirdly, S101, P102, D103 of CDRH3 formed 3 pairs of hydrogen bonds with G545, L546, T547 of SD1, further improving the affinity of the antibody. The conformational change after the unstable depolymerization of SARS-CoV S leads to exposure of more accessible epitopes.

Taken together, PW5-5 and PW5-535 both bind to a hidden epitope in S RBD outside RBM. To further explore whether there is a synergy between PW5-5 and PW5-535, various pseudotyped viruses were tested by neutralization assay. The results showed that the cocktail made of PW5-5 and PW5-535 with non-overlapping epitopes displayed detectable synergy effect (Supplementary Figure 11).

### Prophylactic activity of PW5-5, PW5-535 and PW5-570 in hamster models

To further evaluate the antiviral effects of the isolated neutralizing mAbs, we selected PW5-5, PW5-535, and PW5-570 for further investigation on the golden Syrian hamster model (Figure 7A), based on the results of neutralizing assay with pseudovirus and authentic virus *in vitro*. Since PW5-570 demonstrated the most potent protective effects against SARS-CoV-2 variants from WT to Omicron BA.2, with IC_50_ values less than 0.01 μg/ml, it was selected to evaluate the antiviral effects against SARS-CoV-2 BA.1 on hamsters. Golden Syrian hamsters were treated with one dose of PW5-570 (20 mg/kg) or PBS prophylactically through intraperitoneal injection. At 24 hours post pretreatment, we intranasally inoculated the hamsters with SARS-CoV-2 BA.1, and sacrificed the hamsters at day 2 post virus inoculation. Our results suggested that prophylactic treatment of PW5-570 significantly reduced BA.1 replication in both hamster nasal turbinates and lungs, which was evidenced by the significantly lowered viral RdRp gene copy (Figure 7B) and the undetectable infectious virus titers upon treatment (Figure 7C). We next analyzed the expression of SARS-CoV-2 BA.1 nucleocapsid (N) protein in the harvested hamster lungs. We detected abundant viral N expression in the bronchiole epithelium as well as in the alveolar space in PBS-treated hamster lungs. In striking contrast, viral N gene expression was not detected in PW5-570-treated hamsters (Figure 7D), suggesting the robust anti-SARS-CoV-2 potency of PW5-570.

**Figure 7.**
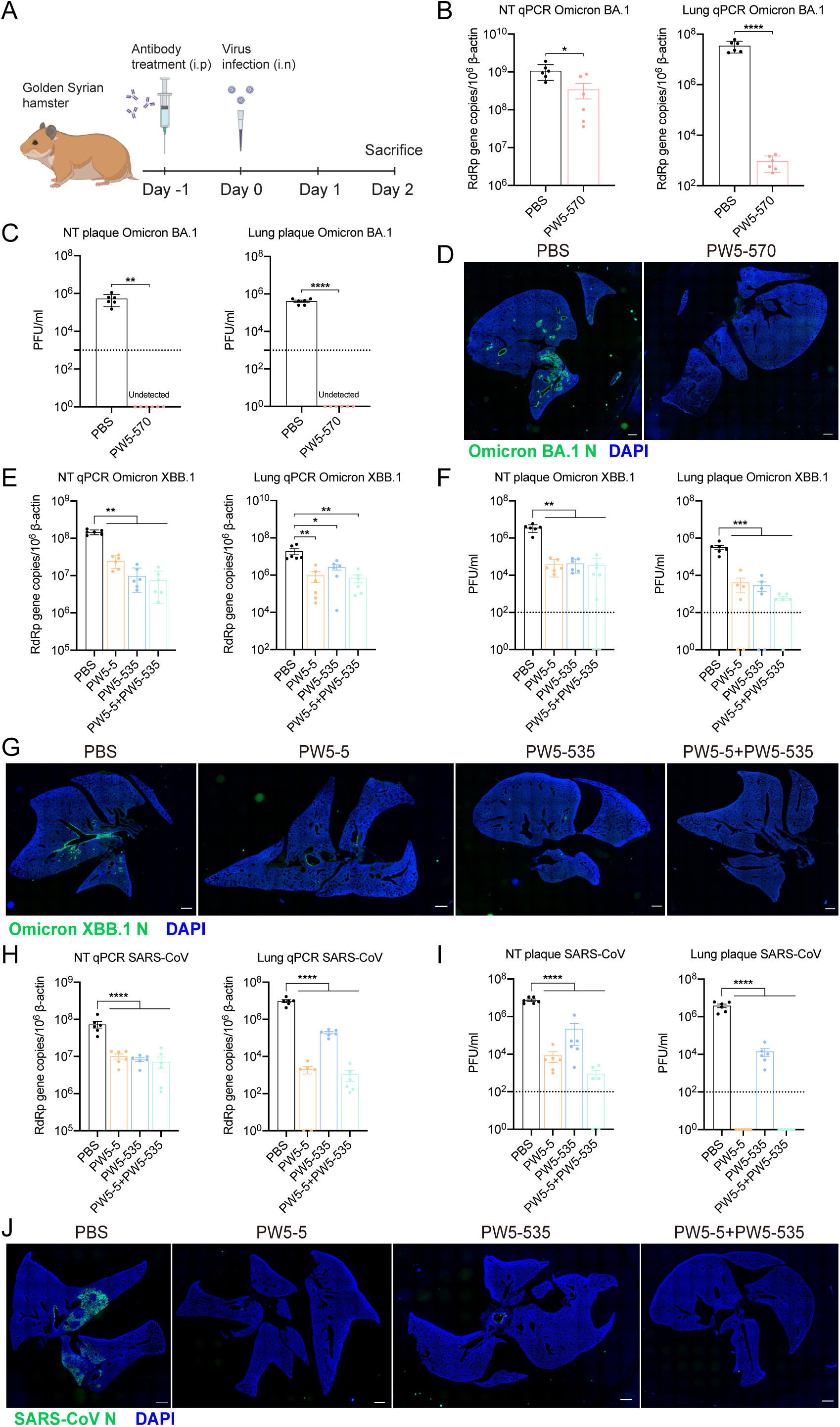
Prophylactic efficacy of PW5-5, PW5-535 and PW5-570 against SARS-CoV, SARS-CoV-2 Omicron BA.1 and XBB.1 in golden Syrian hamsters. **(A)** Schematic of the *in vivo* mAb efficacy test experiment on hamsters. Viral load quantification of Omicron BA.1(**B and C**), XBB.1 **(E and F)** and SARS-CoV **(H and I)** in golden hamster lung and nasal turbinate with or without treatments. Golden hamster lung samples were harvested on day 2 post SARS-CoV-2 Omicron BA.1, XBB.1, or SARS-CoV challenge and homogenized for qRT-PCR analysis and plaque assay titration (n = 6). **(D, G and J)** Immunofluorescence images of the whole section of infected hamster lungs with or without treatments. SARS-CoV-2 nucleocapsid (N) protein was identified with a rabbit anti-SARS-CoV-2-N immune serum (green). Nuclei was identified with DAPI stain (blue). Scale bars in **(D, G and J)** represented 1000 μm for 4-fold magnifications of the objective with 10-fold magnification at the eyepiece. The experiments in **(B-C)** and **(E-F)** and **(H-I)** were repeated two times independently with similar results, respectively. Data represented means and SEMs from the indicated number of biological repeats. Statistical significance between groups in **(B-C)** was determined by unpaired t test and the statistical significance between groups in **(E-F)** and **(H-I)** was determined with one-way ANOVA. *P < 0.05, **P < 0.01, ***P < 0.001, and ****P < 0.0001.

In the meantime, we selected PW5-5 and PW5-535 to investigate their anti-sarbecovirus potential *in vivo* due to their broad-spectrum viral neutralizing activity against sarbecovirus. SARS-CoV-2 XBB.1 and SARS-CoV were selected to be the model sarbecoviruses for investigation. Golden Syrian hamsters were treated with one dose of PW5-5 (20 mg/kg), PW5-535 (20 mg/kg), the combination of PW5-5 and PW5-535 (10 mg/kg for each mAbs) or PBS prophylactically through intraperitoneal injection. The hamsters were inoculated with SARS-CoV-2 XBB.1 or SARS-CoV at 24 hours post treatment. At day 2 post virus inoculation, the hamsters were sacrificed for sample harvest. Our results demonstrated that prophylactic treatment of PW5-5, PW5-535 or the combination of both mAbs all significantly reduced the replication of XBB.1 in the hamster nasal turbinates and lungs, with significantly lowered viral RdRp gene copy (Figure 7E) and infectious virus titers upon treatment (Figure 7F). In parallel, we performed immunofluorescence staining of SARS-CoV-2 N protein in the harvested hamster lungs. As shown in Figure 7G, PW5-5, PW5-535 or PW5-5/PW5-535 dual treatment all attenuated viral N expression in the lungs of infected hamsters when compared with the PBS treatment. Importantly, prophylactic treatment of PW5-5, PW5-535 or the combination of both mAbs also significantly reduced the replication of SARS-CoV in the hamster nasal turbinates and lungs. We observed significantly lowered SARS-CoV RdRp gene copy (Figure 7H) and infectious virus titers upon mAb treatment (Figure 7I). Compared to the abundant expression of viral N protein in the PBS group of SARS-CoV infected hamster lungs, the expression of viral N protein was marginally detected in the lungs with treatment of PW5-5, PW5-535 or the combination of both mAbs (Figure 7J), indicating both PW5-5 and PW5-535 have the potential to neutralize SARS-CoV-2 XBB.1 and SARS-CoV *in vivo*. Overall, our results reveal that the selected neutralizing mAbs of PW5-5 and PW5-535 have broad-spectrum antiviral effects *in vivo*.

## Discussion

In this study, we succeeded in creating multiple antibodies that are effective against a wide range of coronaviruses from a SARS-CoV-2 sequential vaccinated donor with cross-reactive serum neutralizing activity. Among the 86 antibodies generated, PW5-399 and PW5-530 have efficacy against three severe human coronaviruses, although the neutralization potency is not very high. Moreover, PW5-570 displayed the most potent neutralizing activity against all SARS-CoV-2 strains prior to the Omicron BA.5 variant. Of note, PW5-4, PW5-5 and PW5-535 were found to broadly neutralize all sarbecoviruses tested. Furthermore, the binding profiles of PW5-570 demonstrated that it is a class I neutralizing antibody that blocks ACE2 binding through interaction with RBD. Distinct from some other reported mAbs of this class, such as CB6 and REGN10933, PW5-570 retains full activity against all SARS-CoV-2 strains prior to the Omicron BA.5 variant^37, 38^. Sequence alignment of SARS-CoV-2 and SARS-CoV variants, combined with analysis of residues involved in PW5-570, PW5-5 and PW5-535, shows that a large proportion of PW5-570 epitope residues are mutated in the latest variants, and F486V is indeed the key mutation in the S for the antibody evasion (Supplementary Figure 12). In contrast, PW5-5 and PW5-535 bound to more conserved epitopes hidden in RBD with non-overlapping epitopes, most likely belong to class IV mAbs, explaining their breadth in neutralizing different variants. More importantly, PW5-5 and PW5-535 were less affected by Omicron variants, indicating that broadly neutralizing antibodies in this class can also be potential therapeutic antibodies against current VOCs and future emerging variants. In line with two class IV mAbs, GAR20 and AB-3467, PW5-535 not only competed with CR3022 but also blocked binding of ACE2 to SARS-CoV-2 S trimer^39, 40^. Additionally, the cocktail of PW5-5 and PW5-535 displayed detectable synergy, suggesting that using combinations of mAbs with cooperative function would both increase potency and decrease the risk of escape.

It’s important to highlight that the cross-reactive antibodies in this study were identified through a combined approach of single B cell sorting and virtual screening processes. We introduced a pipeline specifically tailored for B-cell epitope and antibody analysis. Initially, the structures of antigens and antibodies were simulated to obtain their 3-D structural features. Then, the potential cross-reactive epitopes were identified by whole-protein surface traversal screening. The antigenicity scores between different mutants were contrasted by conformational epitope BLAST tool, CE-BLAST, which has been previously used to screen the cross-reactive epitopes between dengue virus and zika virus^29^, as well as between SARS-CoV and SARS-CoV-2^41^. Further, for each potential cross-reactive epitope, the ranking of the antibody was obtained by scoring the binding ability based on the micro-environment matching of physic-chemical properties between the epitope regions and CDR regions. Our methodology successfully pinpointed 86 antibodies that showcased potential cross-reactive capabilities. Further experiments showed that a majority of these antibodies exhibited commendable binding ability to three different coronaviruses, and some of them even displayed potent neutralizing activity (Figure 2). This underscores the proficiency of our *in-silico* approach in discerning broad-spectrum antibodies based on structural information.

Structural characterization demonstrated that PW5-570 occupied part of the RBM region like other class I antibodies to prevent receptor attachment, similar to CB6, B38 and C102, the binding region is a concave plane at the top of RBD, which is highly coexisting with the RBM region^31, 42, 43^. The binding region of hACE2 and RBD region is directly blocked by steric hindrance, which is an effective and common neutralizing antibody type, but it lost its neutralizing ability to many Omicron VOCs, as RBM is the region with the most variation of all the variants. In addition, the formation of trimer dimer indicates potential steric hindrance or virions aggregation to neutralize virions. Despite the relatively high sequence conservation of SARS-CoV and SARS-CoV-2, there are differences in antigenicity, and antibodies that can broadly neutralize SARS-CoV-2 VOCs and SARS-CoV are rarely reported. In this study, PW5-5 and PW5-535 had significant neutralizing activities against SARS-CoV-2 VOCs and SARS-CoV. Structural analysis showed that the combination of PW5-5 induced RBD to shift outward and enter the “extra wide open” state, resulting in conformational instability of RBD. This phenomenon is similar to n3130v inducing S trimer to form a “wide-up” state, but the deflection angle is larger, causing S trimer to be more unstable^44^. This conclusion is consistent with our observation that incubation with PW5-5 and Omicron XBB S trimers leads to the gradual decomposition of some trimers into monomers. PW5-535 also prevents ACE2 attachment, and it targets the inner cryptic surface of RBD, partially overlapping with the CR3022 epitopes, but the difference is that CR3022 is highly resistant to Alpha, Beta, Gamma, and Delta, and is not resistant to Omicron. PW5-535 has significant neutralization effect on SARS-CoV-2 VOCs and SARS-CoV. Structural information indicates that PW5-535 binds both the implicit epitopes of RBD and the SD1 frontend adjacent to RBD, and the SD1 sequence is highly conserved in SARS-CoV-2 variants. This epitope may explain the broad-spectrum neutralizing activity of PW5-535. In summary, we analyzed the different binding epitopes and neutralizing mechanisms of PW5-570, PW5-5, and PW5-535 through cryo-EM. The epitopes of PW5-5 and PW5-535 are highly conserved and can be paired with other competitive neutralizing antibodies, indicating potential pairing strategies for cocktail therapy against SARS-CoV-2 and novel emerging coronaviruses.

Apart from the *in vitro* neutralization activity and structural characterization, we also demonstrated that prophylactic treatment with a single dose of PW5-570 fully protected golden Syrian hamsters from infection by the Omicron BA.1 variant, indicating PW5-570 represents a best-in-class anti-SARS-CoV-2 antibody for use in the prophylactic setting. More importantly, we demonstrated that PW5-5 and PW5-535 exhibited excellent *in vivo* prophylactic protection against both the circulating SARS-CoV-2 XBB.1 variant and SARS-CoV in the hamster model, indicating that PW5-5 and PW5-535 has the potential to be developed as pan-sarbecoviruses antivirals for immunotherapy and transmission prevention clinically. To this end, future vaccine design should consider stabilizing the binding interface of the immunogen for interaction with PW5-5 and PW5-535.

There are several limitations in this study to be considered when interpreting the results. To understand the frequency of PW5-5- and PW5-535-like broadly neutralizing mAbs among sequential vaccinees, we need to investigate other responders who show equally potent broadly neutralizing antibody responses. Furthermore, we only tested a single-dose of PW5-5, PW5-535 or PW5-570 for prophylactic efficacy in the hamster model. However, these mAbs should be tested for therapeutic efficacy in the hamster model as well in the future to provide useful information in support of clinical development of these antibodies-like broadly neutralizing antibody. Besides, the Fc-mediated effector functions of these mAbs have not been evaluated. In addition, we did not isolate and characterize mAbs recognizing other regions of S including NTD and S2.

## Materials and methods

### Protein expression and purification

The constructs used for expression of stabilized soluble MERS-CoV, SARS-CoV, SARS-CoV-2 and Omicron BA.1 S2P S trimer proteins were obtained from our previous studies^45^. The mammalian expression plasmid for dimeric soluble ACE2 was purchased from Addgene. SARS-CoV-2 NTD (aa1-305) and RBD (aa319-541) were separately cloned into mammalian expression vector pCMV3 with an 8 × His tag and 2 × Strep-tag II tags at the C terminus. Expi293 cells were used for transient transfection with the suitable S2P stabilized S-expression plasmids or other vectors by using 1 mg/mL polyethylenimine (PEI, Polysciences). The supernatant was harvested and the S trimer was purified using Ni-NTA resin (Invitrogen) in accordance with the manufacturer’s protocol five days after transfection. Prior to use, all proteins were further evaluated for size and purity through SDS-PAGE. *Sorting for S trimer-specific B cells and single-cell B cell receptor sequencing*

Flow cytometry of PBMCs from one healthy donor with 5 times vaccinations were conducted following methods described previously^45^. Briefly, PBMCs were stained with LIVE/DEAD Fixable Yellow Dead Cell Stain Kit (Invitrogen) at ambient temperature for 20 min, followed by washing with RPMI-1640 complete medium and incubation with 10 μg/ml SARS-CoV-2 Omicron BA.1- and MERS-CoV-S trimers with His-tag at 4 °C for 1 h. Afterwards, the cells were washed again and incubated with a cocktail of flow cytometry antibodies, containing CD3 APC-H7 (BD Biosciences), CD19 BV421 (BD Biosciences), CD27 APC (BD Biosciences), and anti-His PE (Biolegend), at 4 °C for 1 h. Stained cells were then washed, resuspended in RPMI-1640 complete medium and sorted for S trimer-specific memory B cells (CD3^−^CD19^+^CD27^+^S trimer^+^ live single lymphocytes). The sorted cells were loaded into the BD Rhapsody single-cell analysis platform, which could effectively capture and separate single cells. The BCR library preparation and quality control were performed according to the manufacturer’s protocol and sequenced on the NovaSeq PE150 platform (Illumina).

### In-silico screening of cross-reactive antibodies

To detect the potential cross-reactive antibodies for SARS-CoV-2 WT, SARS-CoV-2 Omicron and MERS-CoV among all 684 antibodies obtained from BCR sequencing, an *in-silico* pipeline was constructed. It is an integrated computational pipeline that could provide *in-silico* screening of broad-reactive antibodies for a group of antigen mutants, which includes five steps: 1) obtaining the 3-D structures of antigen protein based on four different types of template, 2) mapping the epitope regions on the modelled S protein, including the real crystalized epitopes and simulated epitopes, 3) selecting the broad-spectrum epitope regions by calculating the antigenicity similarity score of each epitope regions across the three viruses through CE-BLAST, 4) generating the antibody structure library by modelling the 3-D structures for all 684 antibodies, 5) calculating the physic-chemical features and structure features for structures from the potential broad-spectrum epitope library and the antibody structure library to perform virtual screening between epitope and antibody. Top-ranking antibodies through the *in-silico* screening were selected for further validation. Detailed information on each step can be found below:

#### Structure Modeling

The 3-D structures of S protein of SARS-CoV-2 WT, Omicron and MERS-CoV were constructed through SWISS-MODEL^46^. Considering the S monomer in the trimer structures contains two different modes, up and down, four different trimer structures including: 1) three RBD up (3-RBD-up), 2) two RBD up and one RBD down (2-RBD-up), 3) one RBD up and two RBD down (1-RBD-up), and 4) three RBD down (3-RBD-down). Structures of four trimer types were illustrated in Supplementary Figure 3. The templates of above four trimer types were illustrated in Supplementary Table 1. Antibody structures were modeled through ABodyBuilder^47^, a fully automated Ab structure prediction server, to generate modeled structures of all 684 paired antibodies in our dataset.

#### Epitope mapping

After structure modeling, the epitopes regions were mapped on the S trimers which involve crystalized real epitopes and simulated virtual epitopes. For crystalized real epitope, we derived 1,349 SARS-CoV-2 S-antibody complexes from SabDab^47^. After removing redundancy (according to sequence similarity of 100%), 1,083 S-antibody complexes which involve 1,083 real epitope regions were remained for further analysis. For virtual epitope simulation, we firstly calculated the accessible surface area (ASA) for each residue in the S protein based on the template structures through Pymol^48^. Then, the surface residues were defined as those residues with ASA over 1 Å^2^. Typically, for each surface residue r, all residues in the S protein within 10 Å were defined as the epitope residues of surface patch P. Here, we generated 10,909 epitopes for both SARS-CoV-2 WT and Omicron S, and 9,608 virtual epitopes for MERS-CoV S.

Cross-reactive epitope prediction: For any real or virtual epitope EP, the cross-protective score between SARS-CoV-2 WT - EP, Omicron - EP, and MERS - EP were calculated through CE-BLAST ^29^. The potential cross-reactive epitopes were defined as those epitope regions that obtain cross-reactive scores above the threshold across all three viruses. By setting the threshold of 0.7 for CE-BLAST score, 132 potential cross-reactive epitopes were detected, including 4 real epitopes and 128 virtual epitopes. After mapping the potential cross-reactive epitope on the templates, each epitope region contains three epitope files (SARS-CoV-2-WT, SARS-CoV-2-Omicron and MERS-CoV).

Antibody screening: For each potential cross-reactive epitope, we screened all the 684 antibodies through the patch model of SEPPA-mAb^30^. Typically, the epitope patch of the cross-reactive epitope and the CDR patch of the antibody were encoded into a group of fingerprints to calculate the binding score between each epitope patch and CDR patch. Then, for each epitope patch, the score of CDR patch against MERS-CoV, SARS-CoV-2 WT and SARS-CoV-2 Omicron were ranked among all 684 antibodies. If one CDR patch can be ranked within the top 5 (<1%) among all 684 mAbs in all three epitope files for one epitope region, this antibody will be considered as the potential cross-reactive antibody for the specific epitope region. Then, 80 antibodies obtained by screening, along with 6 high-frequency antibodies, were selected for further experimental test (Supplementary Table 2).

### Antibody expression and purification

Monoclonal antibodies tested in this study were constructed and produced at Fudan University. For each antibody, variable genes were codon optimized for human cell expression and synthesized by HuaGeneTM (Shanghai, China) into plasmids (gWiz) that encode the constant region of human IgG1 heavy or light chain. Antibodies were expressed in Expi293F (ThermoFisher, A14527) by co-transfection of heavy and light chain expressing plasmids using polyethylenimine (Polyscience) and cells were cultured at 37 °C with shaking at 125 rpm and 8% CO_2_. Supernatants were also collected on day 5 for antibody purification using MabSelectTM PrismA (Cytiva, 17549801) affinity chromatography.

### Production of pseudoviruses

Plasmids encoding the MERS-CoV, SARS-CoV, SARS-CoV-2 and SARS-CoV-2 variants spikes, as well as the spikes with single or combined mutations were synthesized. Expi293F cells were grown to 3×10^6^/mL before transfection with the indicated spike gene using Polyethylenimine. Cells were cultured overnight at 37 °C with 8% CO_2_ and VSV-G pseudo-typed DG-luciferase (G*DG-luciferase, Kerafast) was used to infect the cells in DMEM at a multiplicity of infection of 5 for 4 h before washing the cells with 1xDPBS three times. The next day, the transfection supernatant was collected and clarified by centrifugation at 300g for 10 min. Each viral stock was then incubated with 20% I1 hybridoma (anti-VSV-G; ATCC, CRL-2700) supernatant for 1 h at 37°C to neutralize the contaminating VSV-G pseudotyped DG-luciferase virus before measuring titers and making aliquots to be stored at −80°C.

### Pseudovirus neutralization

Neutralization assays were performed by incubating pseudoviruses with serial dilutions of monoclonal antibodies or sera, and scored by the reduction in luciferase gene expression as described previously^49^. In brief, Vero E6 cells were seeded in a 96-well plate at a concentration of 2×10^4^ cells per well. On the following day, pseudoviruses were incubated with serial dilutions of the test samples in triplicate for 30 min at 37°C. The mixture was added to cultured cells and incubated for an additional 24 h. The luminescence was measured by Luciferase Assay System (Beyotime). IC_50_ was defined as the dilution at which the relative light units were reduced by 50% compared with the virus control wells (virus + cells) after subtraction of the background in the control groups with cells only. The IC_50_ values were calculated using nonlinear regression in GraphPad Prism.

### Viruses and biosafety

SARS-CoV GZ50 (GenBank accession number AY304495) was an archived clinical isolate at Department of Microbiology, HKU. SARS-CoV-2 wild-type (WT) D614G (GISAID: EPL_ISL_497840), B.1.1.7/Alpha (GenBank: OM212469), B.1.351/Beta (GenBank: OM212470), B.1.617.2/Delta (GenBank: OM212471), Omicron BA.1 (EPI_ISL_6841980), Omicron BA.2 (GISAID: EPI_ISL_9845731), Omicron BA.5 (GISAID: EPI_ISL_13777658), Omicron XBB.1 (GISAID: EPI_ISL_15602393) and Omicron BQ.1.1 (GISAID: EPI_ISL_16342297) strains were isolated from the respiratory tract specimens of laboratory-confirmed COVID-19 patients in Hong Kong^50, 51^. SARS-CoV-2 were cultured using Vero-E6-TMPRSS2. SARS-CoV were cultured in Vero-E6 cells. All the viruses were titrated by plaque assays. All experiments with infectious SARS-CoV-2 and SARS-CoV were performed according to the approved standard operating procedures of the Biosafety Level 3 facility at the Department of Microbiology, HKU.

### Authentic SARS-CoV and SARS-CoV-2 variants neutralization

An end-point dilution assay in a 96-well plate format was performed to measure the neutralization activity of select purified mAbs as described previously^45^. In brief, each antibody was serially diluted (5-fold dilutions) starting at 100 μg/ml. Triplicates of each mAb dilution were incubated with indicated live virus at a MOI of 0.1 in DMEM with 7.5% inactivated fetal calf serum for 1 h at 37 °C. After incubation, the virus–antibody mixture was transferred onto a monolayer of Vero-E6 cells grown overnight. The cells were incubated with the mixture for 70 h. CPEs were visually scored for each well in a blinded fashion by two independent observers. The results were then converted into percentage neutralization at a given mAb concentration, and the averages ± S.E.M. were plotted using a five-parameter dose–response curve in GraphPad Prism 8.0.

### Epitope mapping by ELISA

For the epitope binding ELISA, the 50 ng per well of indicated S trimer, 50 ng per well of RBD, and 100 ng per well of NTD were coated onto ELISA plates at 4 °C overnight. The ELISA plates were then blocked with 300 μl blocking buffer (0.5% BSA and 5% Skim Milk) in PBST (0.05% Tween-20 in PBS) at 37 °C for 2 h. Afterwards, purified antibodies were serially diluted using dilution buffer (0.2% BSA and 2% Skim Milk in PBST), incubated at 37 °C for 1 h. Next, 100 μl of 5,000-fold diluted Peroxidase AffiniPure goat anti-human IgG (H+L) antibody (Promega) was added into each well and incubated for 1 h at 37 °C. The plates were washed between each step with PBST in three times. Finally, the TMB substrate (Promega) was added and incubated before the reaction was stopped using 1 M sulfuric acid. Absorbance was measured at 450 nm.

For the competition ELISA, the first antibody at the concentration of 2 μg/mL was coated on ELISA plates and incubated at 4 °C overnight. The ELISA plates were then blocked with 300 μl blocking buffer (0.5% BSA and 5% Skim Milk) in PBST at 37 °C for 2 h. After washing, SARS-CoV-2 S trimer with 2 × Strep-tag II tags was diluted to a final concentration of 2 μg/mL, followed by mixing with 50 μg/mL of the second competition antibodies or PBS blank control. After incubation at 37 °C for 1 h, plates were washed three times with PBST and the 5,000-fold diluted anti-Strep-Tactin HRP conjugate antibody (IBA Life Science) was subsequently added. Then the plates were incubated at 37 °C for 1 h again. The plates were then washed and developed with TMB, and absorbance was read at 450 nm after the reaction was stopped. Competitive percentage of two antibodies was calculated with reference to the PBS blank control.

For the ACE2 competition ELISA, 100 ng of soluble ACE2 protein was immobilized on the plates at 4 °C overnight. The unbound ACE2 was washed away by PBST and then the plates were blocked. After washing, 100 ng of SARS-CoV-2 S trimer with 2 × Strep-tag II tags in 50 μl dilution buffer was added into each well, followed by addition of another 50 μl of serially diluted competitor antibodies and then incubation at 37 °C for 1 h. The ELISA plates were washed three times with PBST and then 100 μl of 5,000-fold diluted anti-Strep-Tactin HRP conjugate antibody (IBA Life Science) was added into each well for another 1 h at 37 °C. The plates were then washed and developed with TMB, and absorbance was read at 450 nm after the reaction was stopped.

### Cryo-EM data collection and processing

The indicated S trimer at 5 mg/mL was mixed with PW5-5, PW5-535 or PW5-570 at 5 mg/mL in a 1:1.2 molar ratio (S trimer/IgG), incubated at 4°C for 1 h, the indicated S trimer-antibody complexes were separated on a Superose 6 Increase 10/300 GL and dilute to 0.7 mg/ml in 20 mM Tris, pH 8.0, 200 mM NaCl. XBB/SARS-CoV S-PW5-535 complexes gradually depolymerize after incubation and then be diluted directly after incubation for freezing samples. A 3 μl complex sample was added to a freshly glow-discharged holey amorphous nickel-titanium alloy film supported by 400-mesh gold grids. Vitrobot IV (FEI, Thermo Fisher Scientific) was used to immerse the sample in liquid ethane and frozen it for 2-s, -3blot force, and 10-s wait time.

Cryo-EM data were collected on a TITAN Krios G4 Transmission electron microscopy (Transmission electron microscopy) operated at 300kV, equipped with a Falcon 4i and a Selectris X Imaging filter (Thermo Fisher Scientific) setting to a slit width of 20 eV. EPU software at 300kV in AFIS mode was used for automated data collection. EER movie stack was collected in super-resolution mode, at a nominal magnification of 105,000x for PW5-535 and PW5-570, and 130,000x for PW5-5, corresponding to a physical pixel size of 1.19 Å and 0.932 Å, dose fractioned to 1737 frames and 1080 frames, respectively. Defocus ranges from −1.0 μm to −3.0 μm, with a total dose of about 50 e-/Å2.

All the data processing was performed using either modules on, or through, RELION v3.1 or cryoSPARC v4.2.1. The movie stacks were binned 2×2, dose weighted, and motion corrected using MotionCorr2 within RELION. Then CTF estimated using Gctf. All micrographs were imported to cryoSPARC for patched CTF-estimation, particle picking and 2D classification. Good particles from selected good micrographs were imported to Relion and then 3D auto refine and 3D classification were performed. The reported resolutions above were based on gold-standard Fourier shell correlation (FSC) 0.143 criterion. All the visualization of maps and handedness correction are performed in UCSF Chimera. Sharpened maps are generated using DeepEMhancer. The resolution was estimated according to gold-standard Fourier shell correlation (FSC) 0.143 criterion. All the data processing procedures are summarized in Supplementary Figure 13–17. The maps were sharpened by DeepEMhancer, and handedness was corrected using UCSF Chimera.

Model building and refinement: SWISS-MODEL and Omicron S-FD01 (PDB:7WOW), SARS-CoV S-CV38-142 (PDBID:7LM9), Omicron BA.1 S-MO1 Fab (PDBID:8H3M) was used for generating initial model, models were fitted into the maps using UCSF Chimera, and further manually adjusted in COOT, followed by several rounds of real space refinement in PHENIX. Model validation was performed using PHENIX. Figures were prepared using UCSF Chimera and UCSF ChimeraX. The statistics of data collection and model refinement are listed in Supplementary Table 3.

### Hamster protection experiment

All experiments in the study complied with the relevant ethical regulations for research. The use of animals followed all relevant ethical regulations and was approved by the Committee on the Use of Live Animals in Teaching and Research of The University of Hong Kong. Male and female golden Syrian hamsters, aged 4-6 weeks old, were obtained from the HKU Centre for Comparative Medicine Research (CCMR). To evaluate the antiviral effects of selected antibodies, hamsters were pre-treated intraperitoneally with one dose of PW5-570 (20 mg/kg), PW5-5 (20 mg/kg), PW5-535 (20 mg/kg), or combination of PW5-5 (10 mg/kg) and PW5-535 (10 mg/kg), respectively, one day before virus inoculation. Then, the hamsters treated with PW5-570 or the vehicles were inoculated with 1×10^4^ PFU SARS-CoV-2 BA.1 pre-diluted in 50 μl PBS intranasally, 24h after antibody pretreatment. Meanwhile, the hamster treated with PW5-5, PW5-535, combination of PW5-5 and PW5-535, or the vehicles were inoculated with 5×10^3^ PFU SARS-CoV or 2×10^4^ PFU SARS-CoV-2 XBB.1 pre-diluted in 50 μl PBS intranasally, respectively. All intranasal treatment in hamsters was performed under intraperitoneal ketamine (100 mg/kg) and xylazine (10 mg/kg) anesthesia. The health status and body weight of hamsters were monitored on a daily basis until the animal was sacrificed or euthanized because of reaching the humane endpoint of the experiment. Hamsters were sacrificed on day 2 post infection. Nasal turbinate (NT) tissues and lung tissues were harvested for immunofluorescence staining, qRT-PCR, or plaque assays as we previously described^52, 53^.

### Immunofluorescence staining

Infected hamster lungs were fixed for 24 h in 10% formalin. The fixed samples were washed with 70% ethanol and embedded in paraffin by TP1020 Leica semi-enclosed benchtop tissue processor (Leica biosystems, Buffalo Grove, IL, USA) and sectioned with microtome (Thermo Fisher Scientific). Sectioned samples were dewaxed by serially diluted xylene, ethanol, and double-distilled water in sequence. To retrieve the antigens, the sectioned samples were boiled with antigen unmasking solution (H-3300, Vector Laboratories) at 85 °C for 90s, followed by Sudan black B and 1% BSA blocking for 30 min, respectively. The in-house rabbit anti-SARS-CoV-2-N immune serum or in-house rabbit anti-SARS-CoV immune serum was used as the primary antibodies to stain the N protein of SARS-CoV-2 or SARS-CoV accordingly by incubation at 4 °C overnight. The secondary antibody, goat anti-rabbit IgG (H+L) cross-adsorbed secondary antibody (A-11008), was purchased from Thermo Fisher Scientific. The antifade mounting medium with DAPI (H-1200, Vector Laboratories, Burlingame, CA, USA) was used for mounting and DAPI staining. Images were acquired by using Nikon Ti2-E Widefield Microscope (Japan, Tokyo).

### Plaque assays

Vero-E6-TMPRSS2 cells were seeded in 12-well plates at day 0. The harvested supernatant samples were serially diluted by 10-fold and inoculated to the cells for 2h at 37°C. After inoculation, the cells were washed with PBS in three times, and covered with 2% agarose/PBS mixed with DMEM/ 2% FBS at 1:1 ratio. The cells were fixed with 4% paraformaldehyde after 72 h incubation. Fixed samples were stained with 0.5% crystal violet in 25% ethanol/distilled water for plaque visualization.

## Acknowledgement

We thank Center of Cryo-Electron Microscopy, Fudan University for the supports on cryo-EM data collection. This study was supported by funding from the National Natural Science Foundation of China (32270142 to P.W.; 31900483 to T.Q.; 32070657 to Z.C.), National Key R&D Program of China (2019YFA0905900 to Z.C.), Shanghai Rising-Star Program (22QA1408800 to P.W.), Shanghai Sailing Program (19YF1441100 to T.Q.), the Program of Science and Technology Cooperation with Hong Kong, Macao and Taiwan (23410760500 to P.W.), the Ministry of Science and Technology of China (2021YFC2302500 to L.S.) and R&D Program of Guangzhou Laboratory (SRPG22-003 to L.S.). This study was also supported by Collaborative Research Fund (HKU C7103-22G to H.C.), Theme-Based Research Scheme (T11-709/21-N to H.C.), the Research Grants Council of the HKSAR; the Health and Medical Research Fund (COVID1903010-Project 14 to H.C.), the Food and Health Bureau, the Government of the HKSAR; and Emergency COVID-19 grant (2021YFC0866100 to H.C.) from Major Projects on Public Security under the National Key Research and Development Program of China. Pengfei Wang acknowledges support from Open Research Fund of State Key Laboratory of Genetic Engineering, Fudan University (No. SKLGE-2304) and Xiaomi Young Talents Program. Xiaoyu Zhao acknowledges support from International Postdoctoral Exchange Fellowship Program (Talent-Introduction Program, YJ20220079).

## Author contributions

P.W., L.S., H.C. and Z.C. conceived and supervised the project. X.Z., R.Q., J.L., X.W., C.Li, Y.Cui, C.Z, M.L., Y.Chen, G.C. and W.G. conducted the biological experiments. T.Q., T.M., and Y.W. conducted the bioinformatics analysis. X.H., C.Luo, and C.Y. performed authentic neutralization assays and animal experiments. Q.M. and Y.W. performed the collection of Cryo-EM data and structure determination. J.C. collected the PBMC and serum. X.Z., T.Q., X.H., Q.M., Z.H., W.Z., Z.C., H.C., L.S. and P.W. analyzed the results and wrote the manuscript. All the authors reviewed, commented, and approved the manuscript.

---

**Supplementary Table 1.** Templates of S trimer used in the modeling.

| Trimer type | PDB ID | Resolution | Target |
| --- | --- | --- | --- |
| 3-RBD-up | 7CAK | 3.58Å | SARS-CoV-2 |
| 2-RBD-up-1-RDB-down | 7WOB | 3.25 Å |  |
| 1-RBD-up-2-RDB-down | 7WPA | 2.77Å |  |
| 3-RBD-down | 7OAN | 3.0Å |  |
| 3-RBD-up | 5X59 | 3.7Å | MERS-CoV |
| 2-RBD-up-1-RDB-down | 5X5C | 4.10Å |  |
| 1-RBD-up-2-RDB-down | 5X5F | 4.2Å |  |
| 3-RBD-down | 6Q04 | 2.8Å |  |

**Supplementary Table 2.**
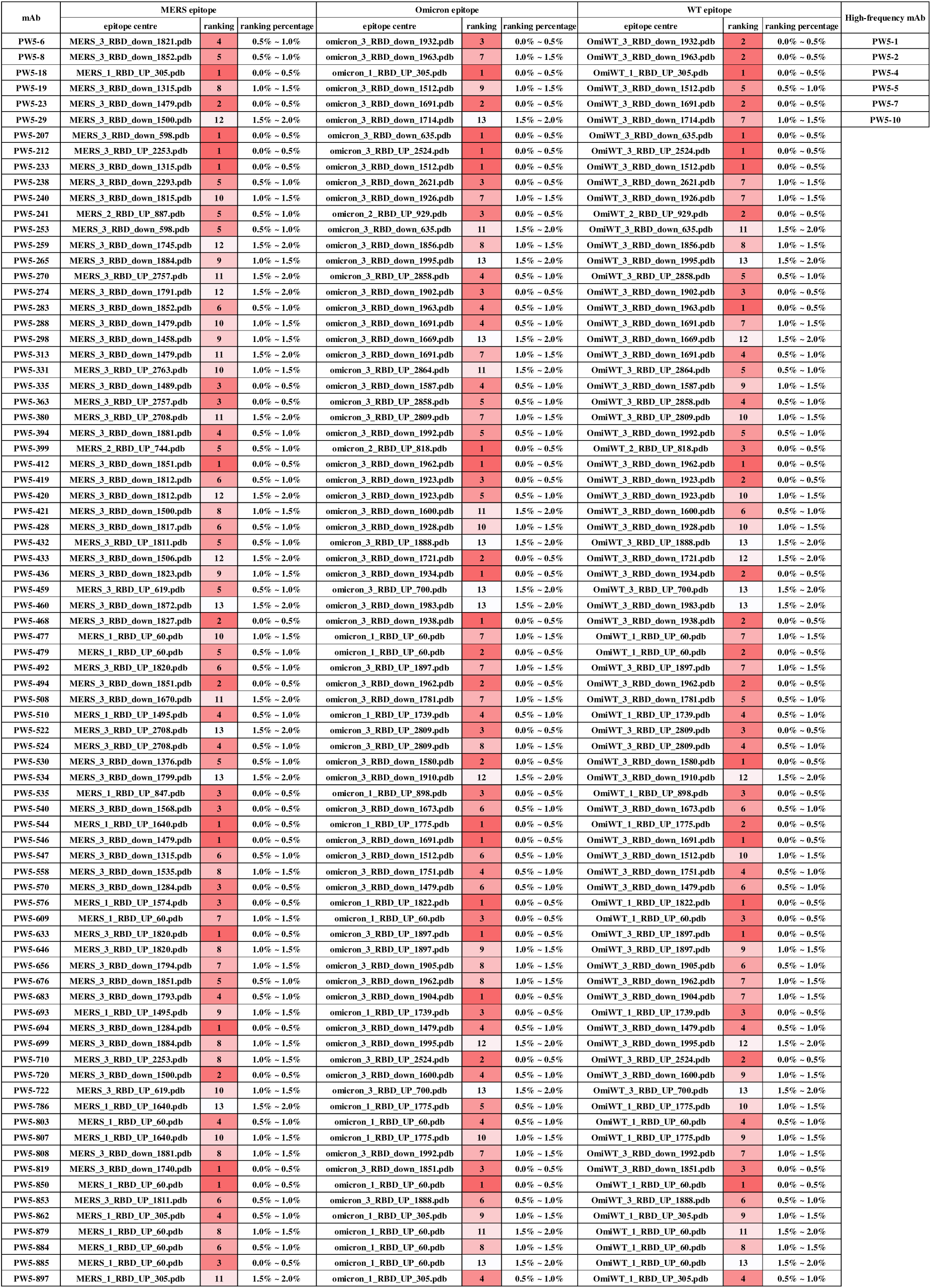
Selected antibodies by in-silico screening.

**Supplementary Table 3.**
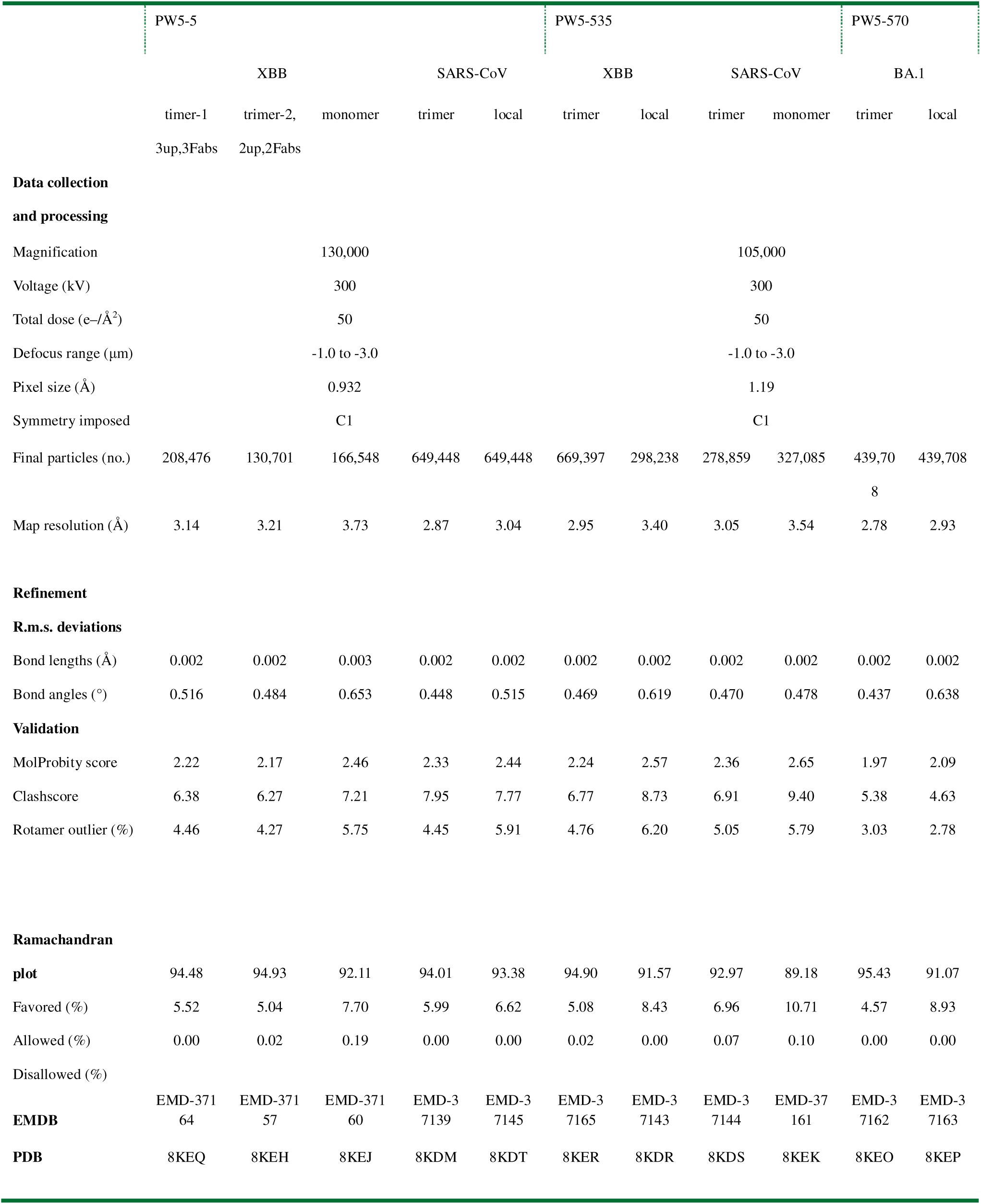
Cryo-EM data collection and refinement statistics.

## Supplementary Figures

**Supplementary Figure 1.**
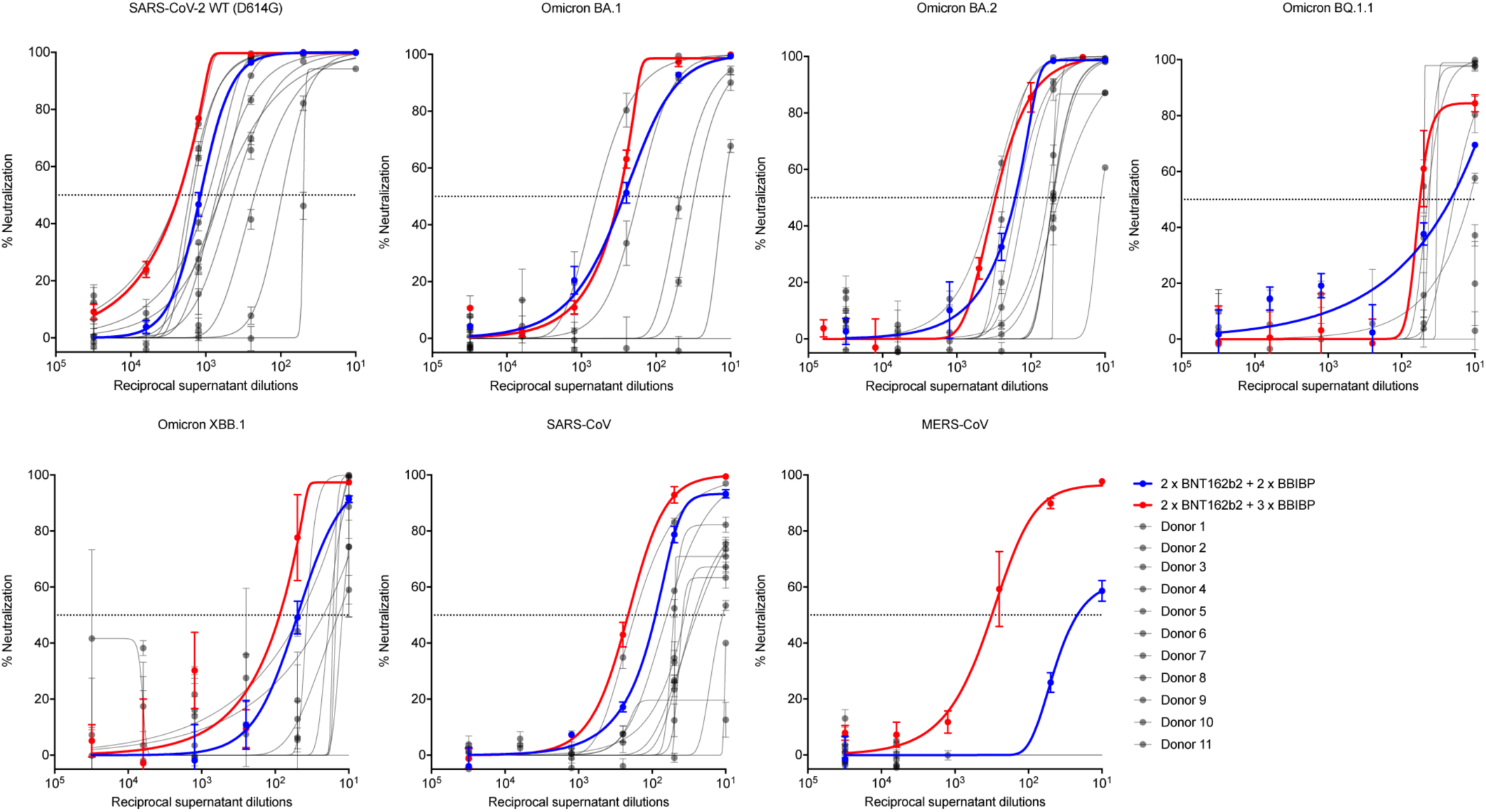
Neutralization curves for sera collected from a healthy volunteer after the fourth- (2 x BNT162b2 + 2 x BBIBP-CorV) or the fifth-vaccination dose (2 x BNT162b2 + 3 x BBIBP-CorV), or from individuals vaccinated with three-dose BBIBP-CorV only (n = 11) against indicated coronavirus. The data are representative of one of at least three independent experiments and are presented as the mean ± SEM.

**Supplementary Figure 2.**
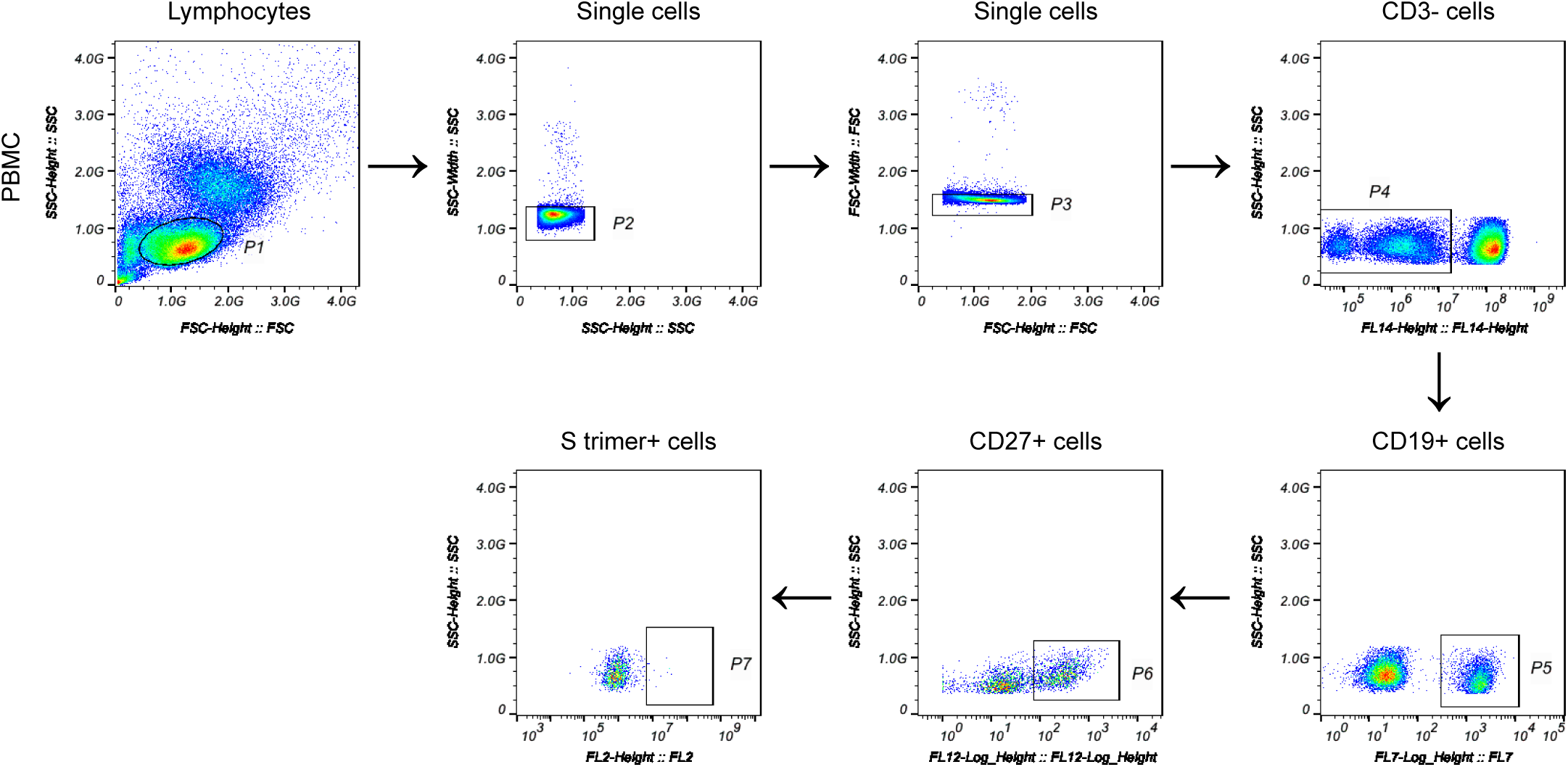
Sorting strategy for the isolation of MERS-CoV or SARS-CoV-2 Omicron S trimer–specific memory B cells using flow cytometry. The sorting primarily focused on live memory B lymphocytes that were CD3^−^, CD19^+^, and CD27^+^. The final step focused on those cells that bind to the MERS-CoV or SARS-CoV-2 Omicron S trimer.

**Supplementary Figure 3.**
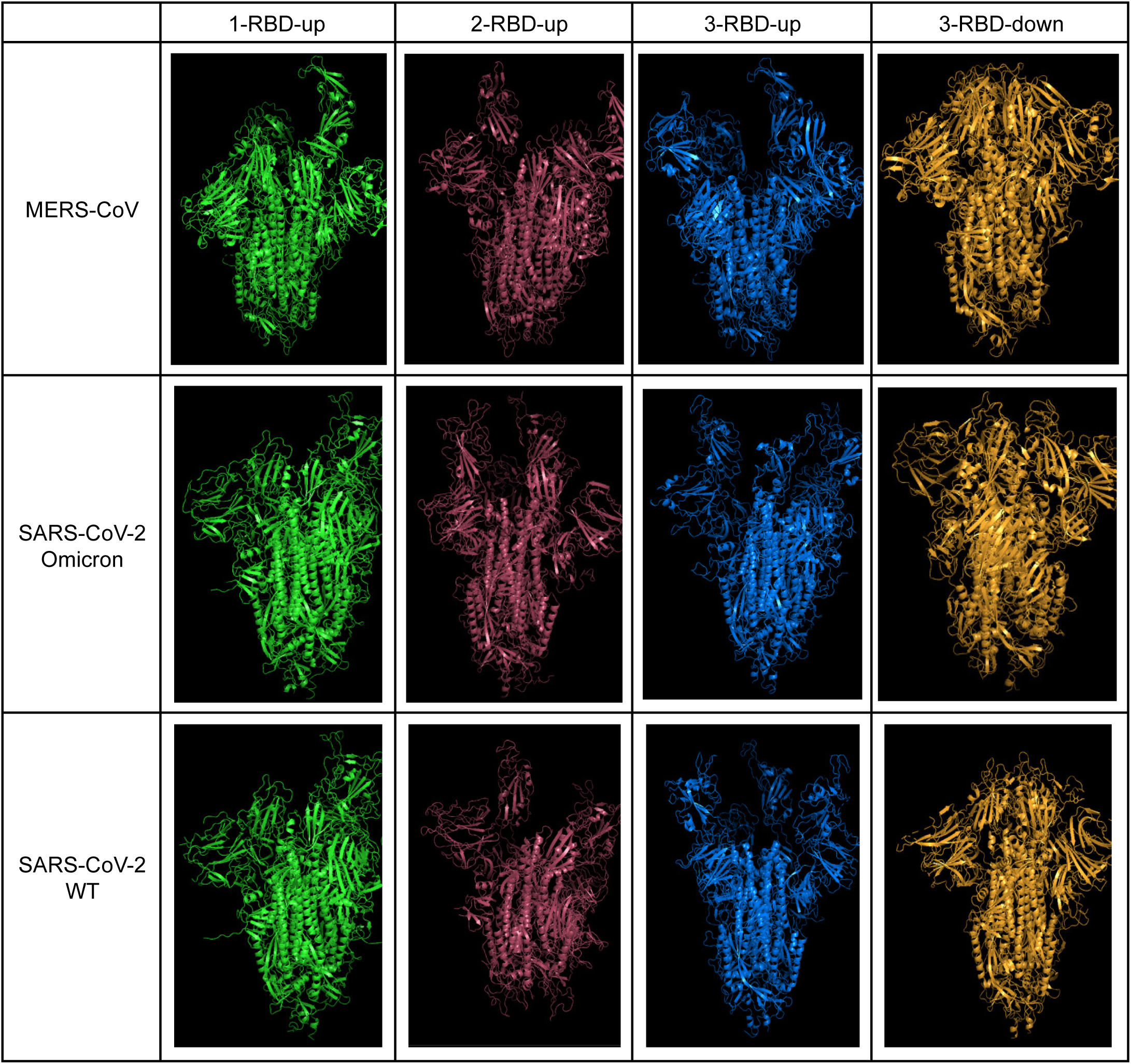
Structure illustration of four S trimer types of SARS-CoV-2 WT, Omicron and MERS-CoV.

**Supplementary Figure 4.**
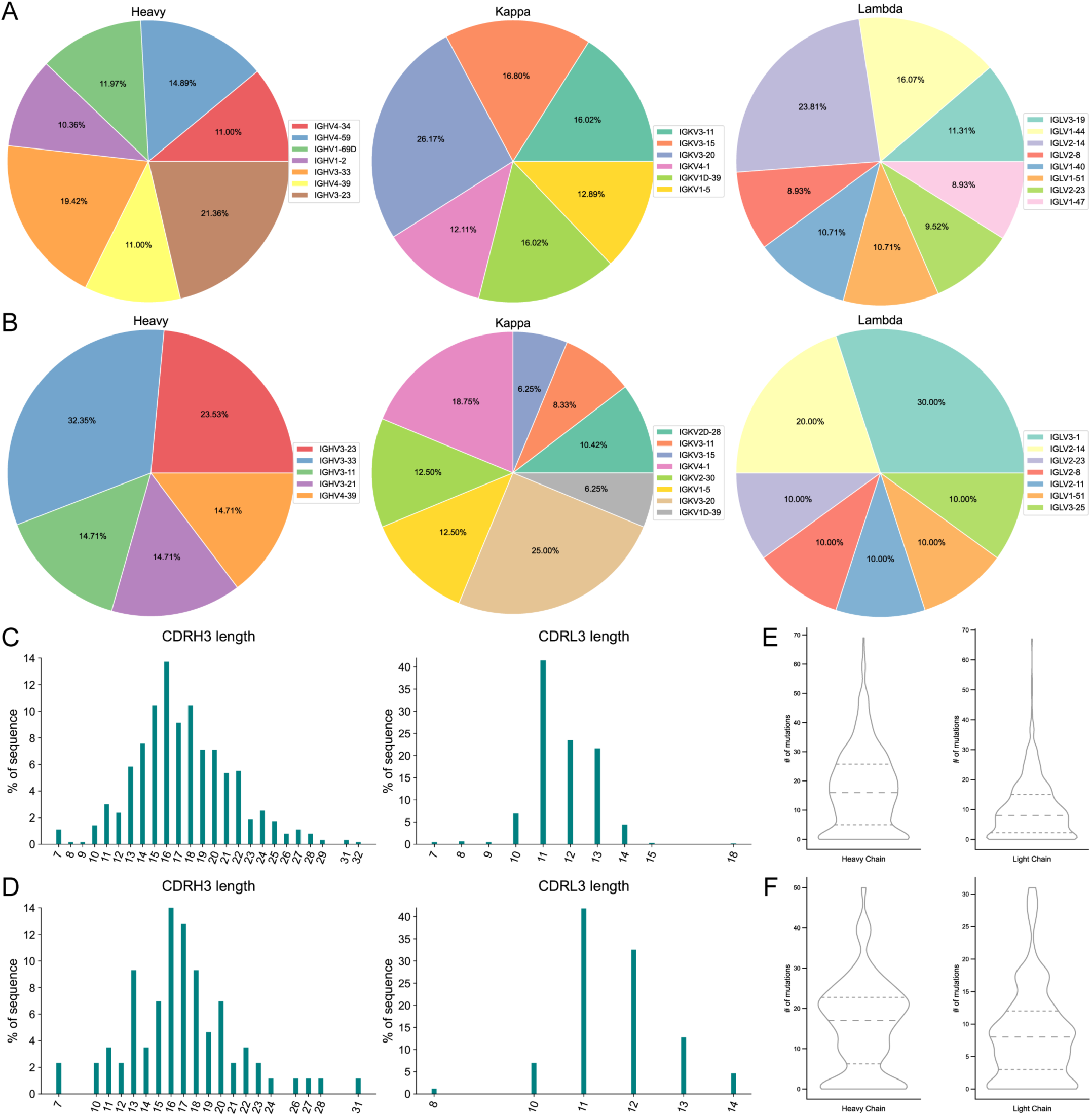
The V gene usage was assigned for paired heavy, kappa and lambda chains recovered from all the 684 sorted S trimer-specific mAbs **(A)** and the 86 mAbs further selected by bioinformatic analysis **(B)**. Percentages are shown on graph for V chains that make up more than 5% of the total for each sort. The CDR3 length distribution for the heavy and light chains of the 684 mAbs **(C)** and the 86 mAbs **(D)**. The number of nucleotide mutations in heavy and light chains of the 684 mAbs **(E)** and the 86 mAbs **(F)**.

**Supplementary Figure 5.**
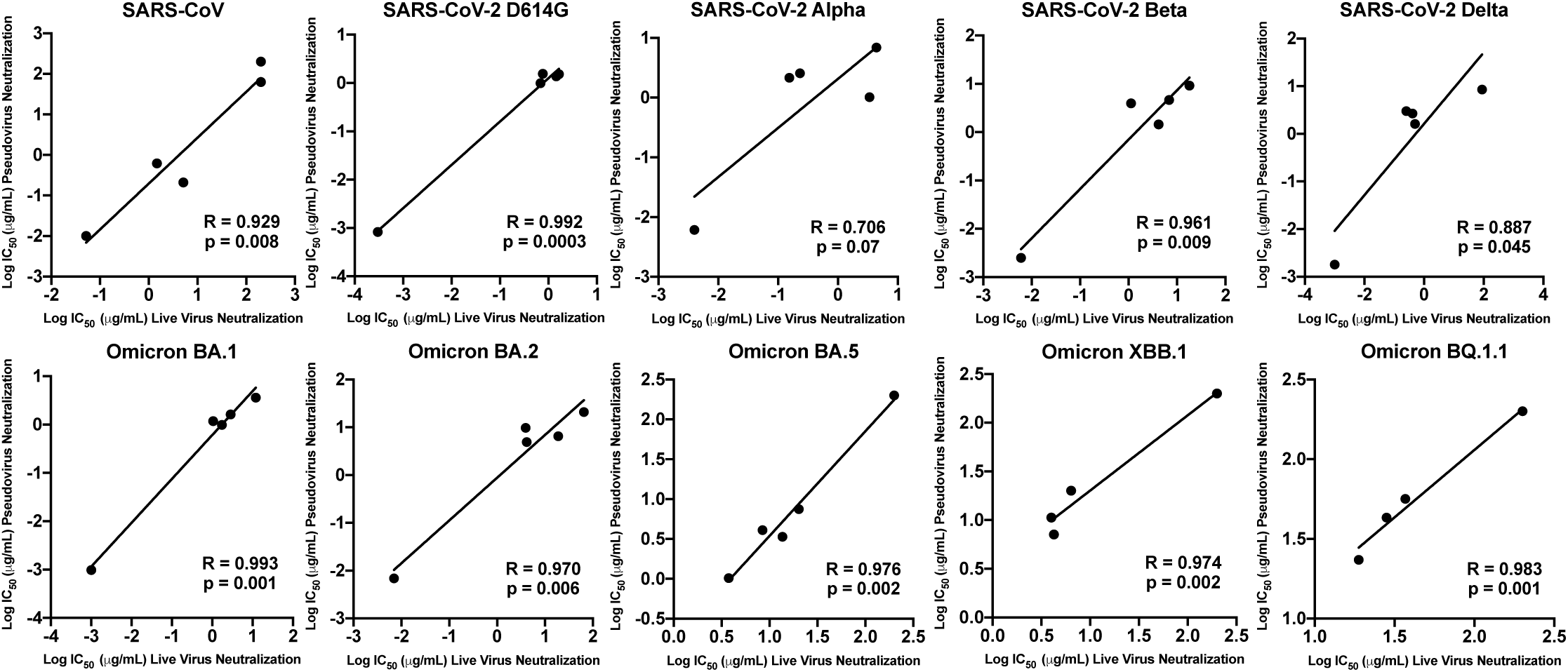
Correlation of live virus neutralization titers versus pseudovirus neutralization titers against SARS-CoV, SARS-CoV-2 and SARS-CoV-2 variants. The Pearson correlation coefficient (R) and the p value were calculated using GraphPad Prism. Experiments were performed in triplicates for all mAbs tested.

**Supplementary Figure 6.**
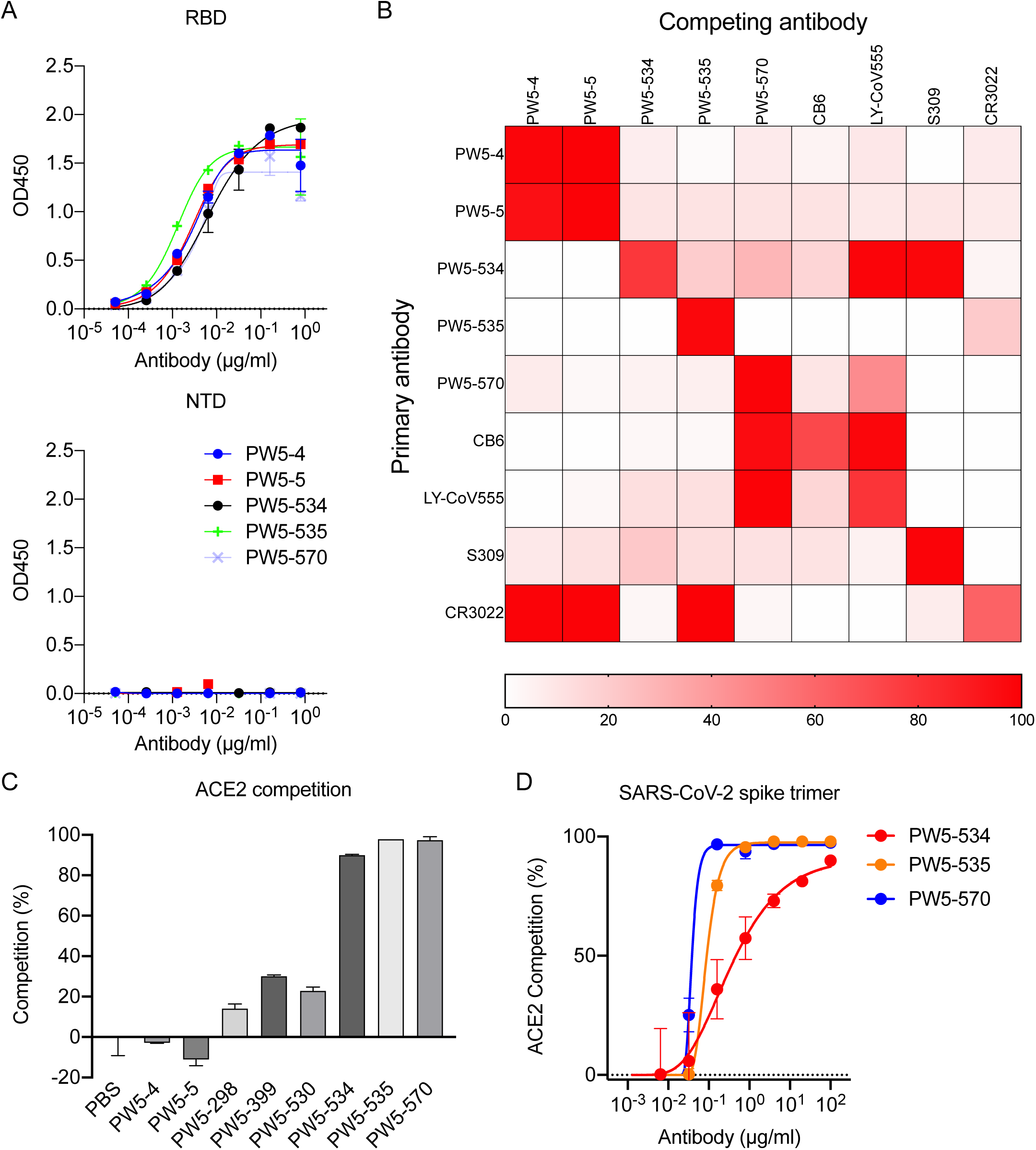
Binding properties of the selected mAbs. **(A)** Binding profiles of 5 purified potent neutralizing mAbs against the RBD (up), and NTD (down) of SARS-CoV-2. **(B)** Epitope mapping of the selected mAbs using a competition ELISA. A heatmap was used to show the competition percentages between two antibodies, as compared with CB6, LY-CoV555, S309 and CR3022, whose epitopes have been well defined. **(C)** Antibodies competitively blocked the ACE2 binding to SARS-CoV-2 S trimer as measured by ELISA. Recombinant human ACE2 protein and phosphate buffer solution (PBS) was used as controls. **(D)** PW5-534, PW5-535, and PW5-570 blocked the ACE2 binding to SARS-CoV-2 S-trimer in a dose dependent manner. The data are representative of one of at least three independent experiments and are presented as the mean ± SD.

**Supplementary Figure 7.**
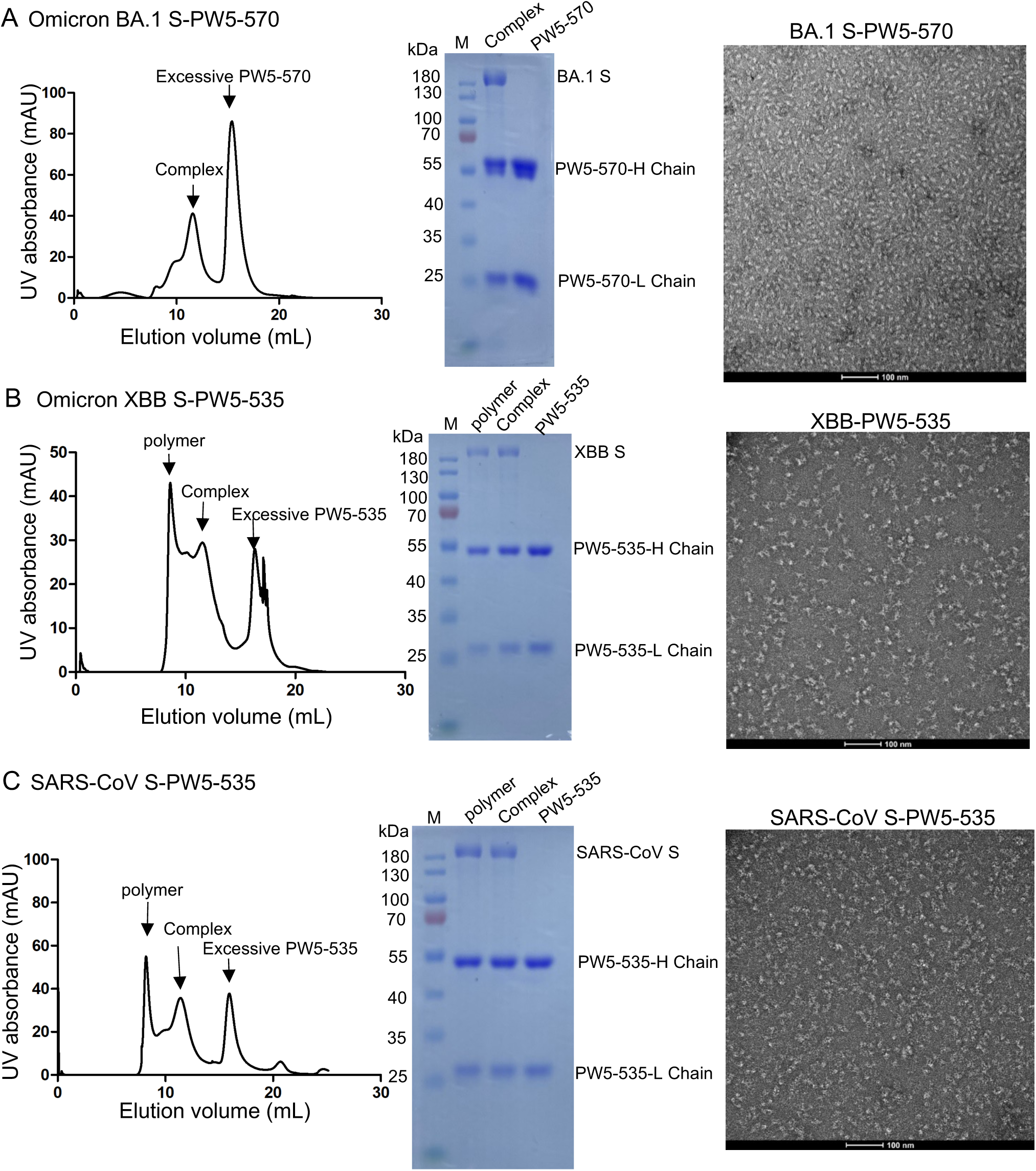
Sample purification of BA.1 S-PW5-570, XBB S-PW5-535 and SARS-CoV S-PW5-535 complex. (**A**) Purification and negative stain images of BA.1 S-PW5-570 complex. (**B-C**) Purification and negative stain images of XBB S-PW5-535 complex and SARS-CoV S-PW5-535 complex.

**Supplementary Figure 8.**
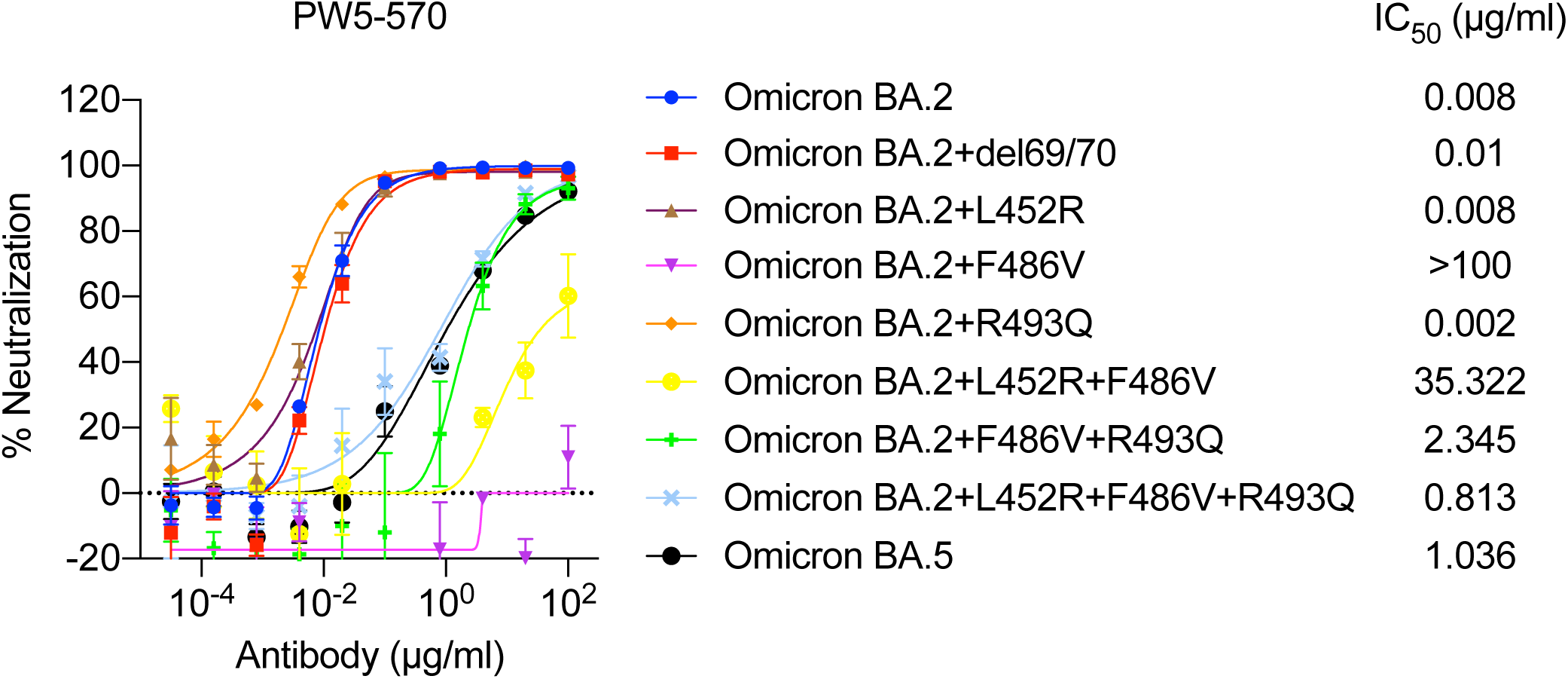
Key mutations conferring PW5-570 antibody resistance. Neutralization of Omicron BA.2 and Omicron BA.5, as well as the Omicron BA.2-based VSVs with single or combined mutations by PW5-570. The data are representative of one of at least three independent experiments and are presented as the mean ± SEM.

**Supplementary Figure 9.**
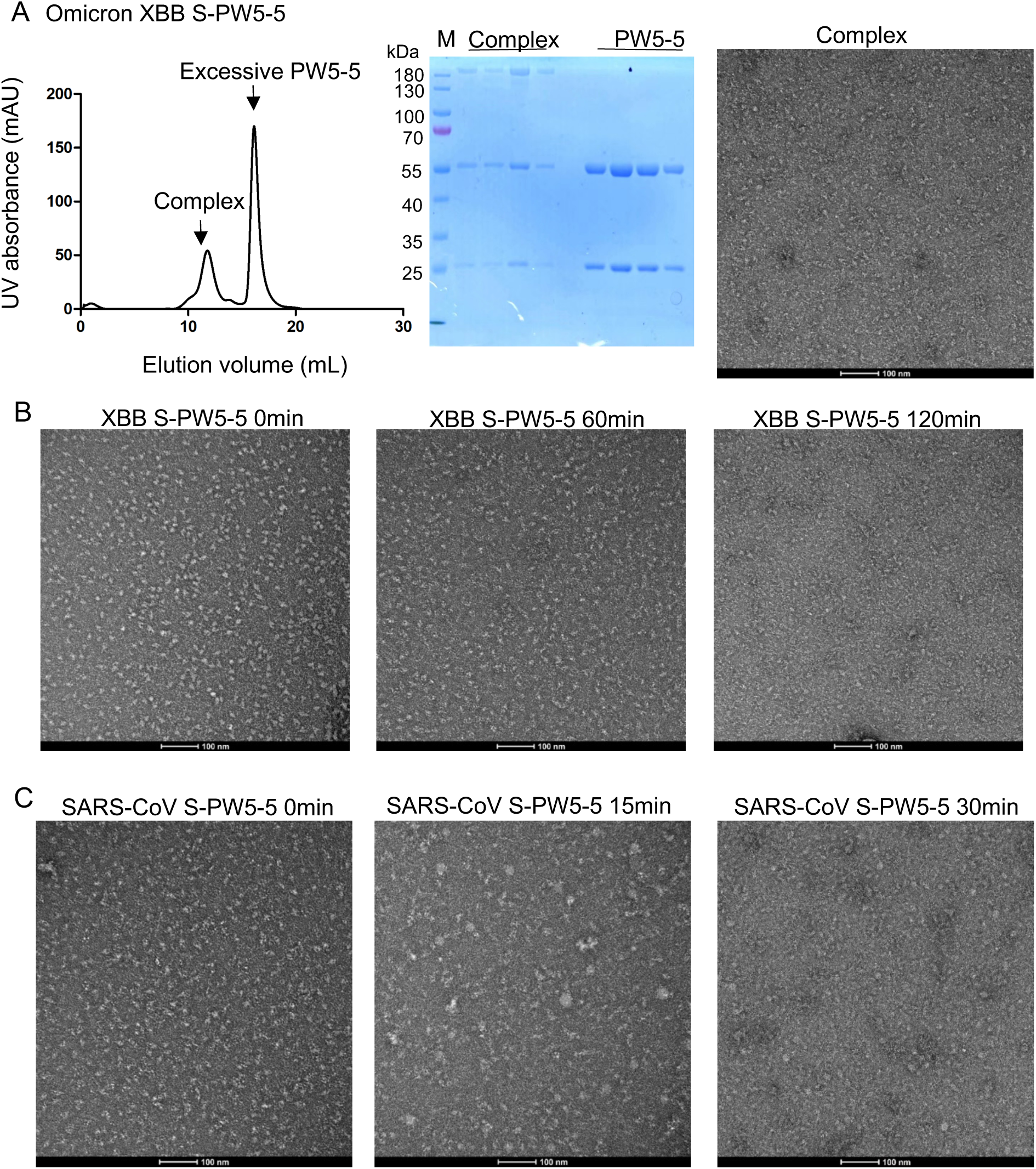
Sample purification of XBB S-PW5-5 and SARS-CoV S-PW5-5 complex. (**A**) Purification of XBB S-PW5-5 complex. The gel-filtration curve showed that PW5-5 and S protein can form complex. (**B-C**) Negative stain images of XBB S-PW5-5 complex and SARS-CoV S-PW5-5 complex, showing that incubating S with PW5-5 induced S trimer disassembling.

**Supplementary Figure 10.**
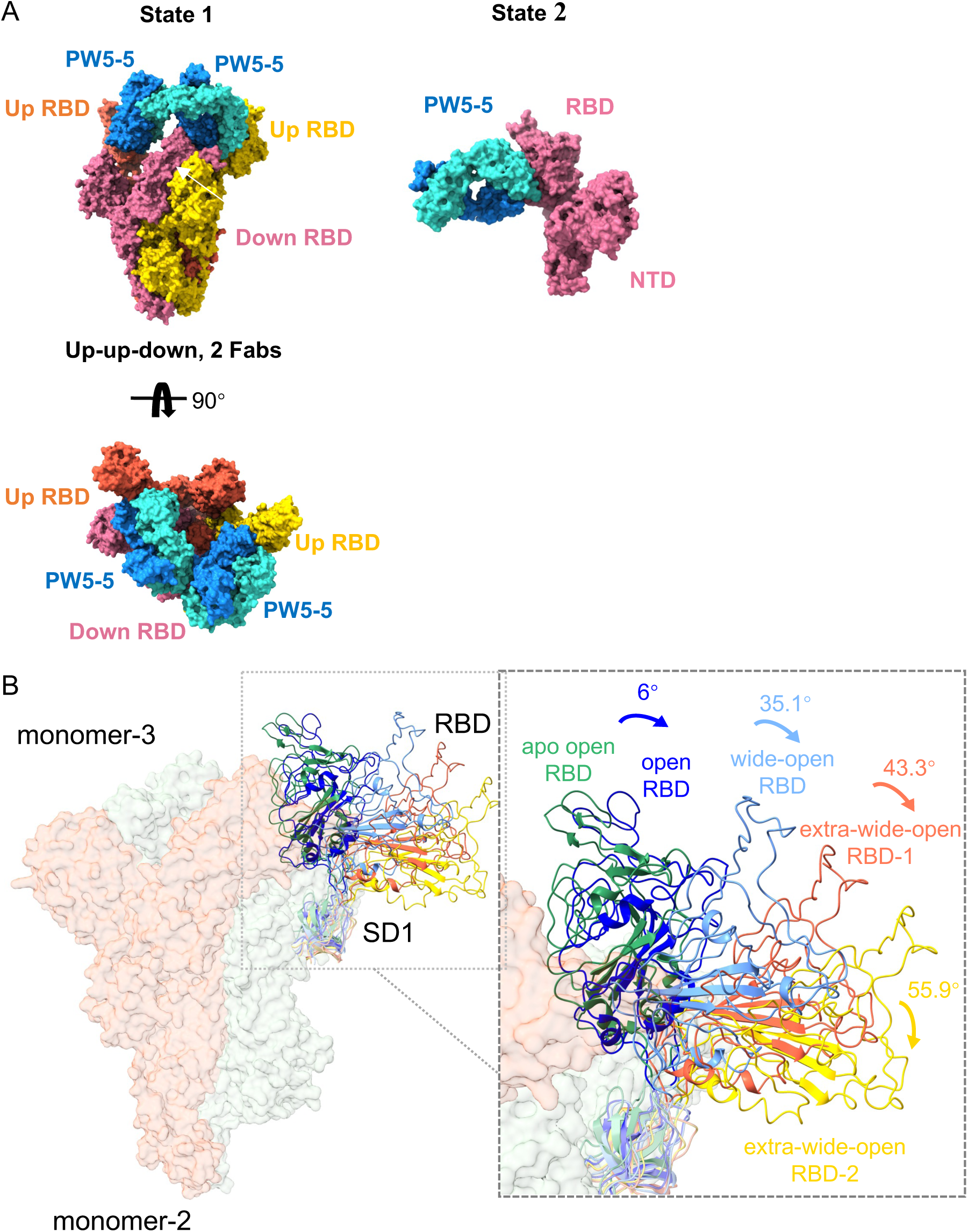
**(A)** Cryo-EM structures of the SARS-CoV S in complex with antibody PW5-5. **(B)** PW5-5 induce obvious movement of RBD. Green: apo state up RBD (PDBID: 7WVN). Dark blue: up RBD bound with bispecific antibody FD01 (PDBID: 7WOQ). Light blue: up RBD bound with nanobody bn03 (7WHK). Orange: up-RBD 1 in 3-up state bound with PW5-5. Gold: up-RBD 2 in 3-up state bound with PW5-5.

**Supplementary Figure 11.**
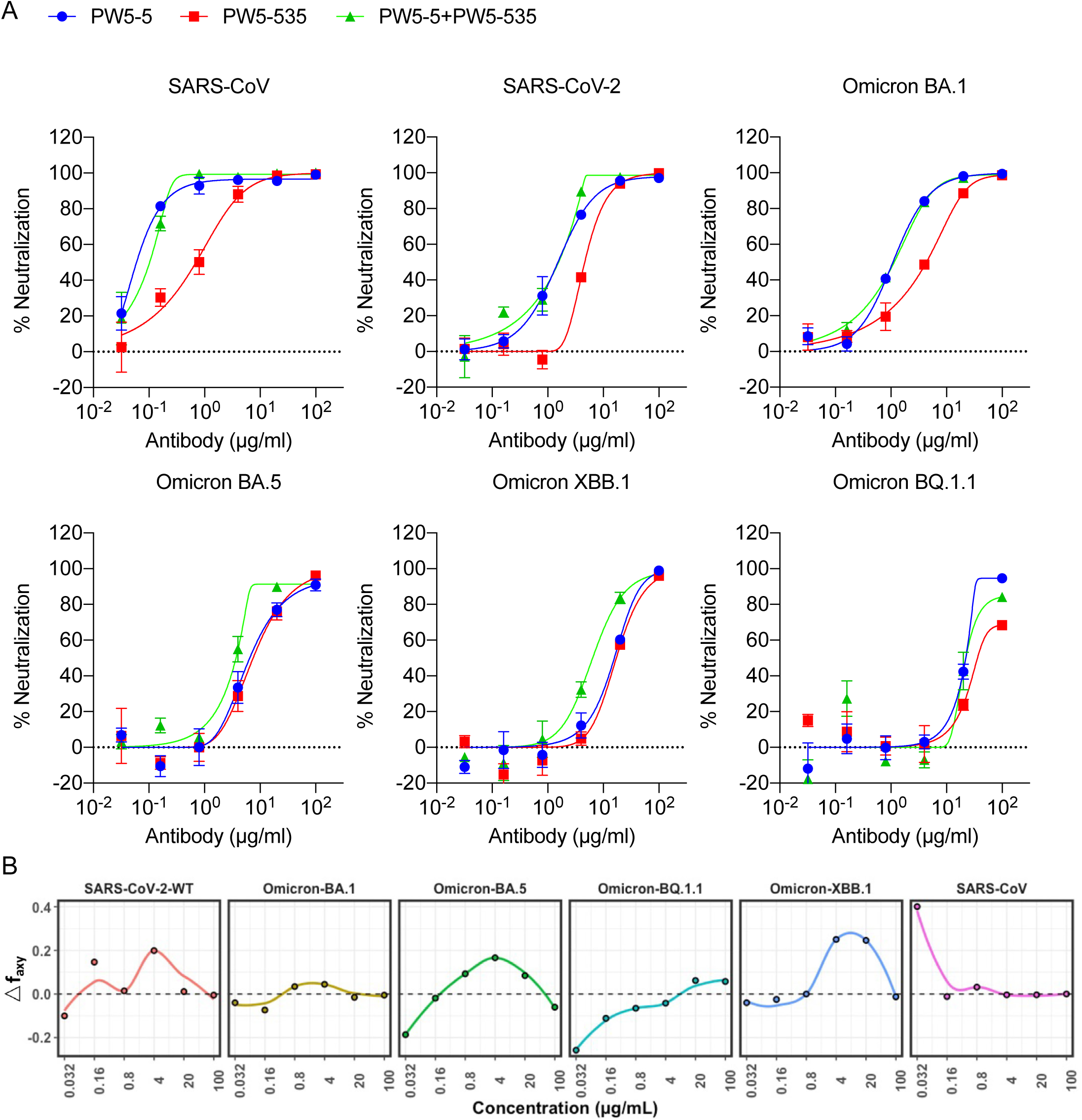
Neutralization synergy effect by PW5-5 and PW5-535. **(A)** Neutralization of SARS-CoV, SARS-CoV-2, and current Omicron subvariants by PW5-5, PW5-535 and their combination, respectively. **(B)** The Bliss independence model was utilized for calculation of synergy for PW5-5 and PW5-535 combinations. Synergy is defined as Δf_axy_ > 0, while Δf_axy_ < 0 indicates antagonism. The data are representative of one of at least three independent experiments and are presented as the mean ± SEM.

**Supplementary Figure 12.**
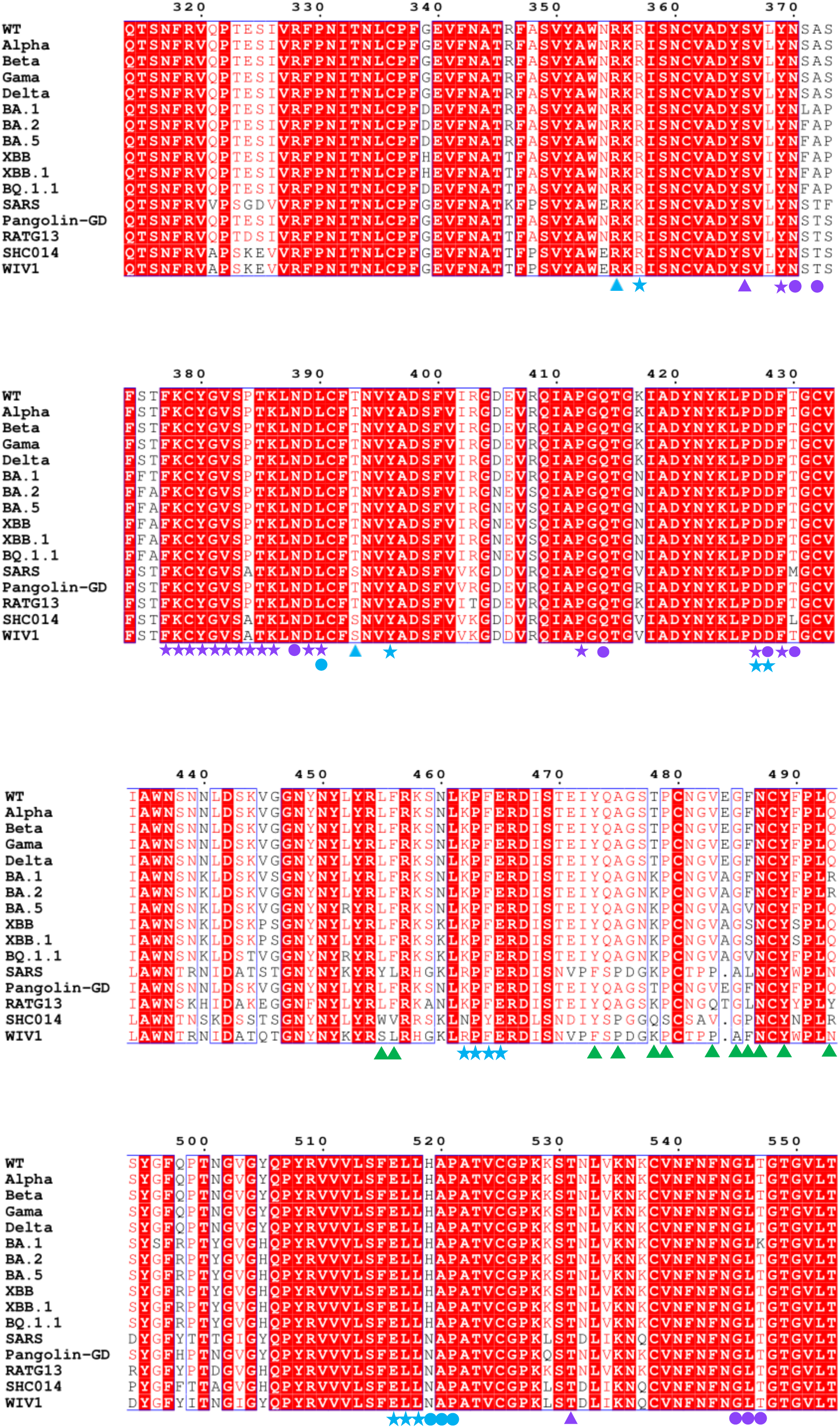
Sequence alignment of SARS-CoV-2 variants, SARS-CoV and other sarbecoviruses. Conserved amino acids are highlighted as red. Residues involved in PW5-570, PW5-5 and PW5-535 binding are marked in green, blue and purple, respectively. Residues involved only in XBB S binding are marked in triangles, only in SARS-CoV S binding are marked in circles and both in XBB and SARS-CoV S binding are marked in stars.

**Supplementary Figure 13.**
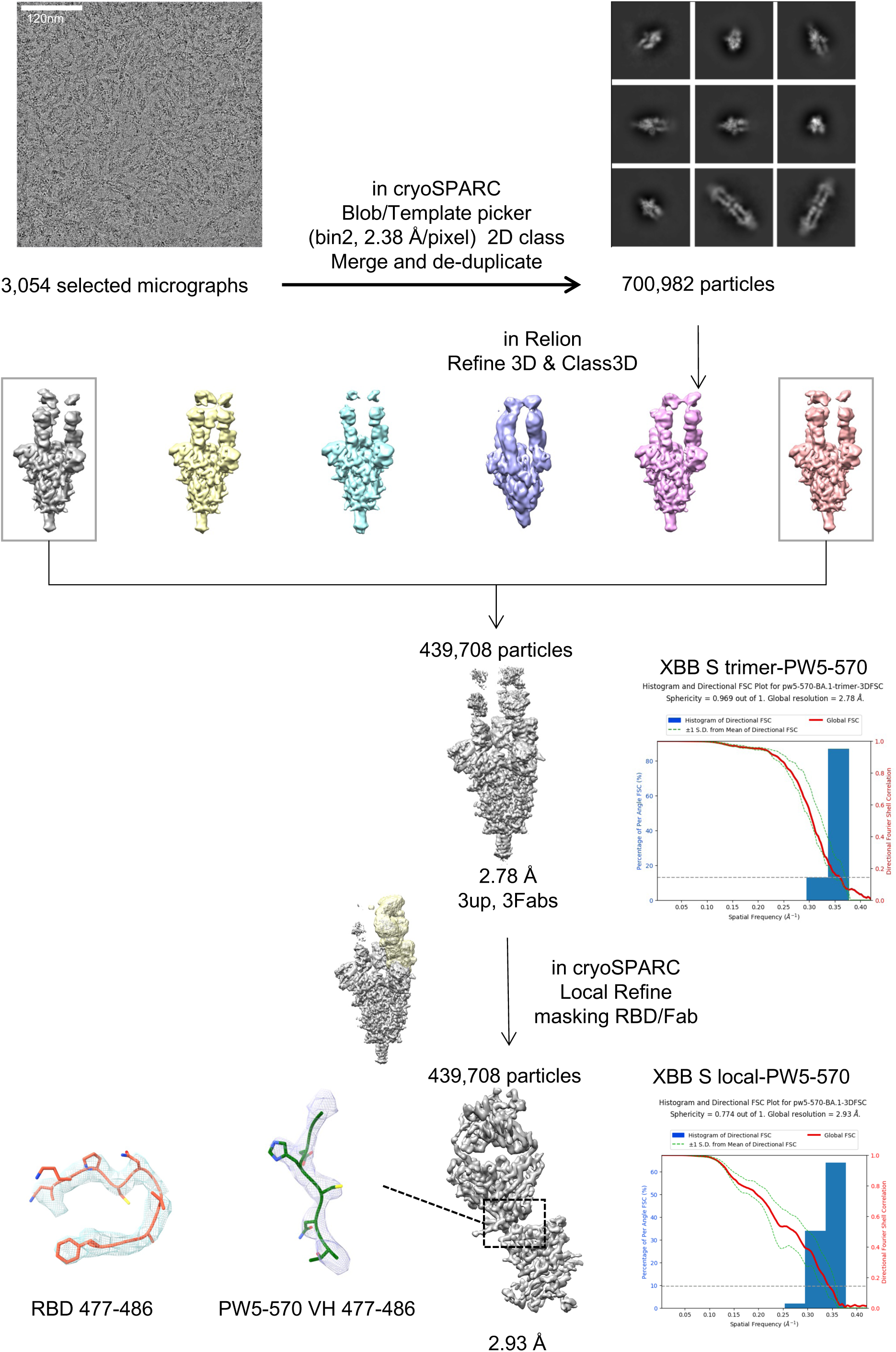
Data processing flowchart of BA.1-S-PW5-570 complex.

**Supplementary Figure 14.**
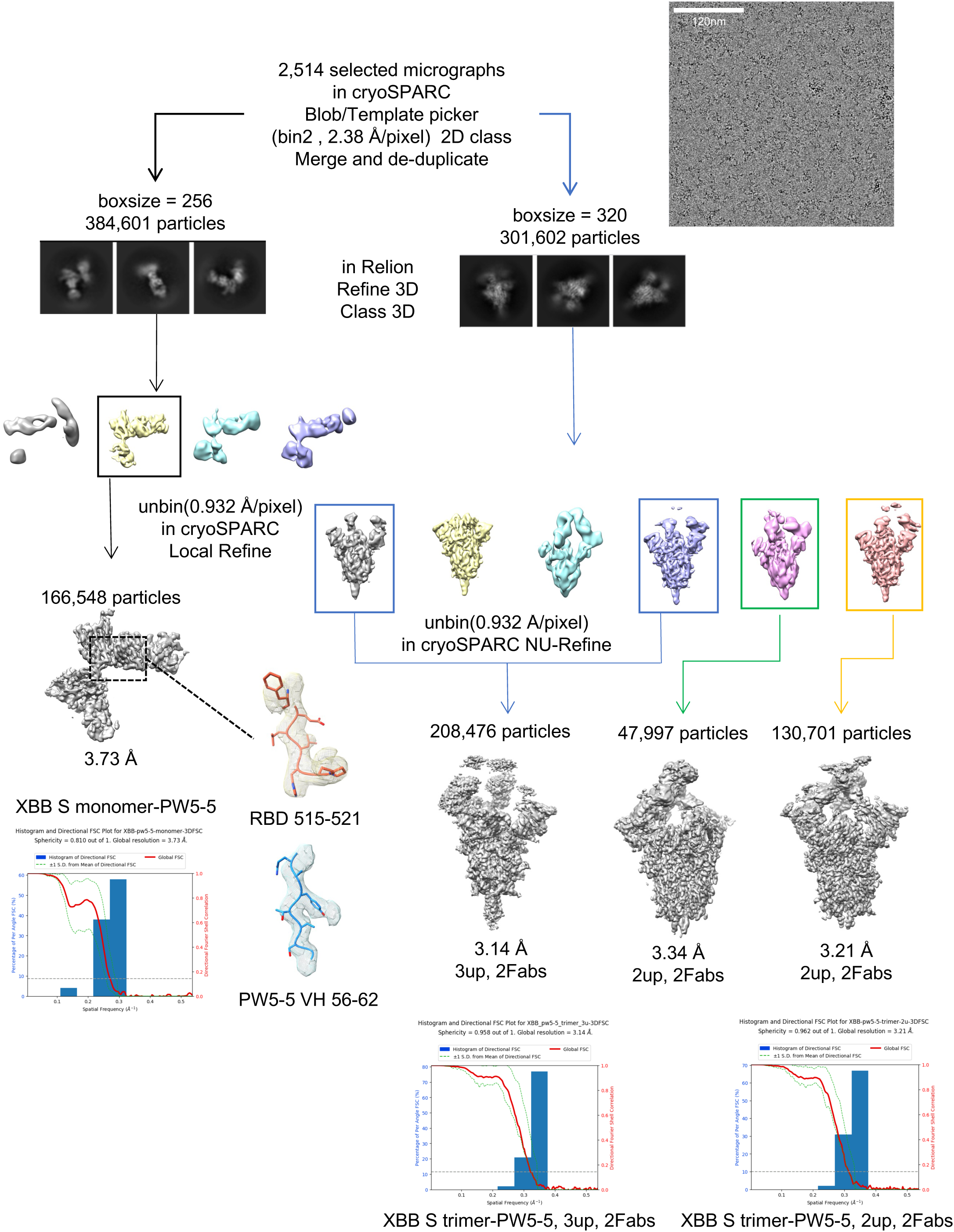
Data processing flowchart of XBB-S-PW5-5 complex.

**Supplementary Figure 15.**
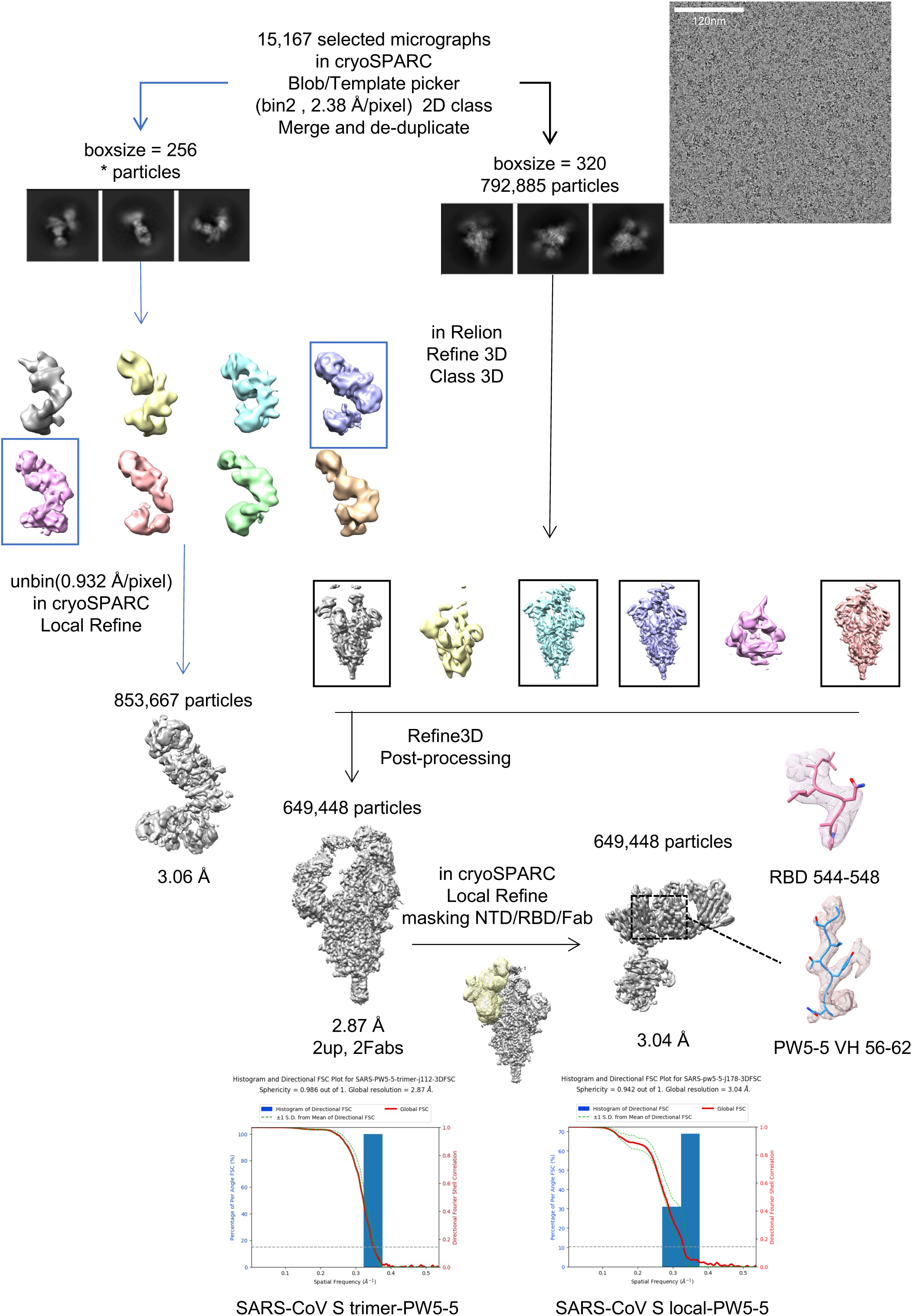
Data processing flowchart of SARS-CoV-S-PW5-5 complex.

**Supplementary Figure 16.**
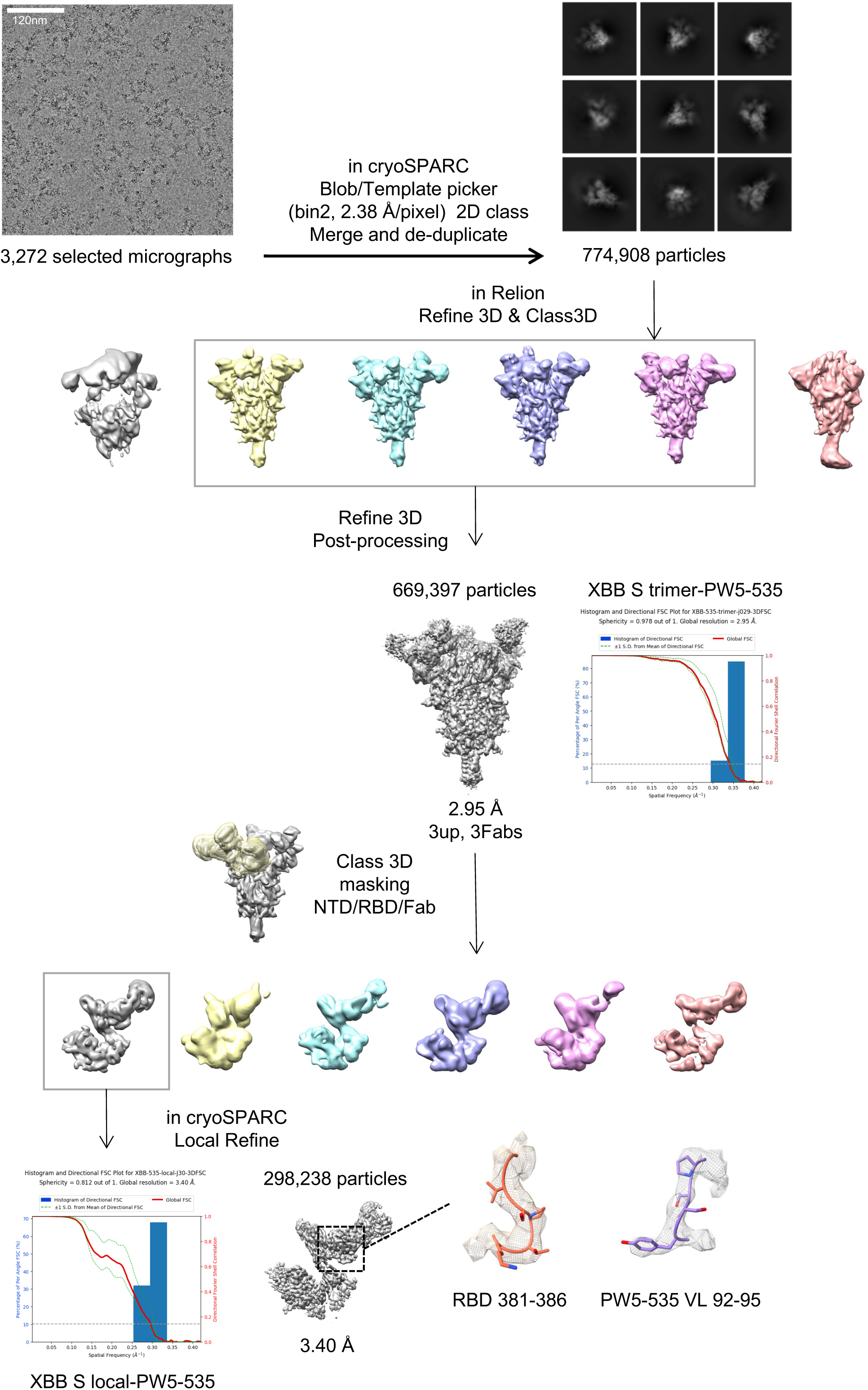
Data processing flowchart of XBB-S-PW5-535 complex.

**Supplementary Figure 17.**
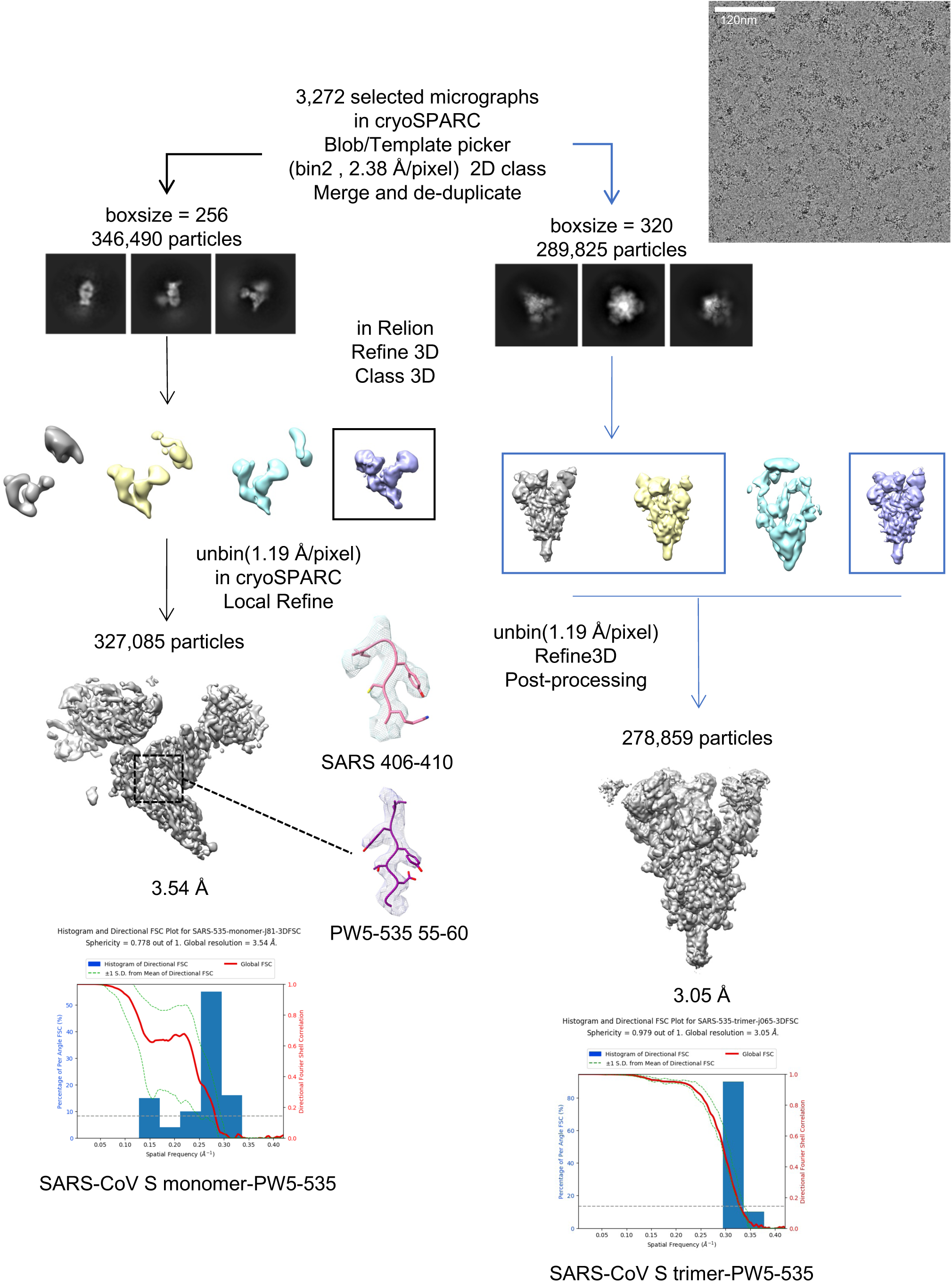
Data processing flowchart of SARS-CoV-S-PW5-535 complex.

